# Structural basis of rotavirus RNA chaperone displacement and RNA annealing

**DOI:** 10.1101/2020.10.26.354233

**Authors:** Jack P. K. Bravo, Kira Bartnik, Luca Venditti, Emma H. Gail, Chen Davidovich, Don C Lamb, Roman Tuma, Antonio N. Calabrese, Alexander Borodavka

**Affiliations:** Department of Biochemistry, University of Cambridge, 8 Tennis Court Road, Cambridge, CB2 1QW, UK; Astbury Centre for Structural Molecular Biology, School of Molecular and Cellular Biology, Faculty of Biological Sciences, University of Leeds, Leeds, UK; Department of Chemistry, Center for NanoScience (CeNS), Nanosystems Initiative Munich (NIM) and Centre for Integrated Protein Science Munich (CiPSM), Ludwig Maximilian University of Munich, Munich, Germany; Department of Biochemistry and Molecular Biology, Biomedicine Discovery Institute, Faculty of Medicine, Nursing and Health Sciences, Monash University, Clayton, Victoria, Australia; EMBL-Australia and the ARC Centre of Excellence in Advanced Molecular Imaging, Clayton, Victoria, Australia; Faculty of Science, University of South Bohemia, Ceske Budejovice, Czech Republic

## Abstract

Rotavirus genomes are distributed between eleven distinct RNA molecules, all of which must be selectively co-packaged during virus assembly. This likely occurs through sequence-specific RNA interactions facilitated by the RNA chaperone NSP2. Here, we report that NSP2 auto-regulates its chaperone activity through its C-terminal region (CTR) that promotes RNA-RNA interactions by limiting its helix-unwinding activity. Unexpectedly, structural proteomics data revealed that the CTR does not directly interact with RNA, whilst accelerating RNA release from NSP2. Cryo-electron microscopy reconstructions of an NSP2-RNA complex reveal a highly conserved acidic patch poised towards RNA. Virus replication was abrogated by charge-disrupting mutations within the acidic patch but completely restored by charge-preserving mutations. Mechanistic similarities between NSP2 and the unrelated bacterial RNA chaperone Hfq suggest that accelerating RNA dissociation whilst promoting inter-molecular RNA interactions may be a widespread strategy of RNA chaperone recycling.

## Introduction

Selective incorporation of viral genomes into nascent virions is essential for virus replication. This process is highly challenging for RNA viruses with multi-segmented genomes (including rotaviruses) since they must coordinate the selection and assembly of 11 distinct RNAs (*1, 2*). Despite these challenges, rotaviruses achieve highly efficient, selective and stoichiometric genome assembly through a series of redundant, sequence-specific intermolecular RNA-RNA contacts (*3*). While RNA-RNA interactions may underpin genome assembly, the remarkable selectivity of these interactions is determined by a complex network of RNA-protein interactions (*4*). In rotaviruses (RV), the viral RNA chaperone protein NSP2 facilitates sequence-specific intersegment RNA-RNA interactions to ensure robust assembly of complete viral genomes (*5, 6*).

NSP2 is a multivalent, non-specific RNA chaperone with high nM affinity for ssRNA (*4, 7*). This allows it to both act as a matchmaker of intermolecular duplexes and limit transient, non-specific RNA-RNA interactions (*4, 8*). This creates a mechanistic conundrum, as NSP2 has to balance helix unwinding and RNA annealing in order to achieve accurate and stoichiometric assembly of distinct RNAs. As such, this viral RNA chaperone plays an absolutely critical role in RV replication (*9, 10*).

Previous mutational studies of NSP2 have been hindered by the lack of a robust reverse genetics system (*11*–*13*). As such, the only region of NSP2 experimentally demonstrated as essential for virus replication to date is the C-terminal region (CTR) (residues 295 – 316) (**Fig. 1a**) (*14*–*16*). We have recently demonstrated that C-terminally truncated NSP2 (NSP2-ΔC) has significantly reduced RNA annealing activity in vitro (*17*). This collectively suggests that the CTR is required for the RNA chaperone activity of NSP2, although its exact functional role(s) remained unclear.

**Fig 1.**
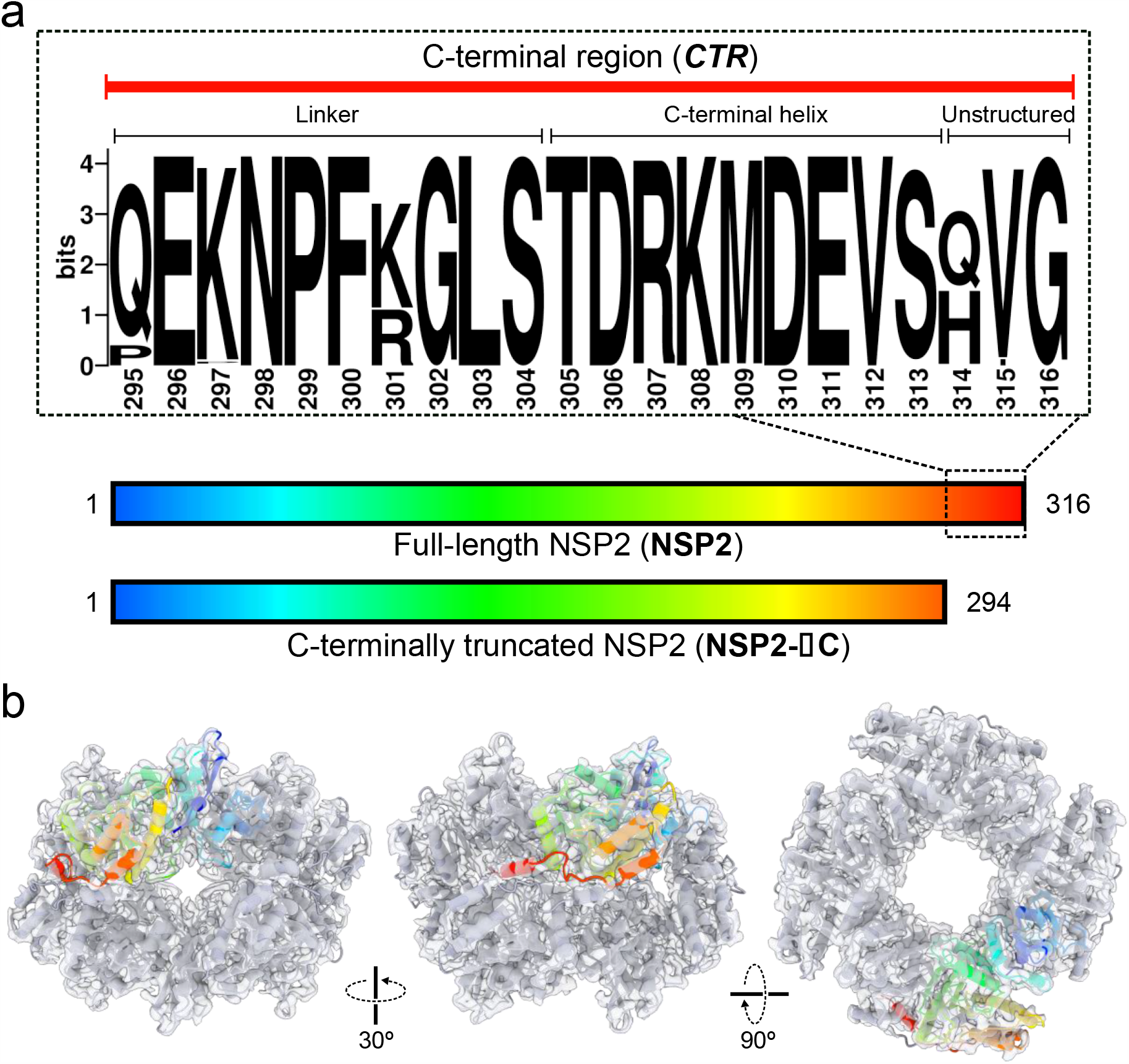
Structure and conservation of NSP2 CTR. **(a)** Constructs of full-length NSP2 (NSP2) and C-terminally truncated NSP2 (NSP2-ΔC, residues 1 – 294) used in this study. An expanded, annotated sequence logo of the NSP2 CTR is shown, which consists of an unstructured, flexible linker region (residues 295 – 304) and a single alpha helix (C-terminal helix, CTH, residues 305 – 313). Downstream residues (314 – 316) are non-essential for viral replication. **(b)** A 3.9 Å resolution cryo-EM reconstruction of the octameric NSP2 apoprotein (grey transparent surface) with associated model (cartoon). A single monomer is highlighted and color coded according to sequence position shown in (a).

Here, we used a single-molecule fluorescence spectroscopy approach to decouple the RNA annealing and RNA unwinding activities of full-length NSP2 and NSP2-ΔC. While NSP2-ΔC exhibits a reduced capacity to promote RNA-RNA interactions, it possesses enhanced RNA unwinding activity. To resolve these paradoxical observations, we determined cryo-EM structures of NSP2 and an NSP2-ribonucleoprotein (RNP) complex at global resolutions of 3.9 Å and 3.1 Å, respectively. In the RNP structure, the RNA density localized to surface-exposed positively-charged grooves, with no evidence of the CTR interacting with the RNA. To directly map the RNA-binding surfaces of NSP2, we employed complementary structural proteomics tools that revealed that all RNA-protein contacts required for non-specific, high-affinity RNA recognition were outside the CTR.

Furthermore, our data show that, while CTR does not directly interact with RNA, it contains a conserved acidic patch that is poised towards bound RNA. Mechanistically, we demonstrate that that the CTR promotes RNA release, indicating that the CTR is required for preventing the formation of a highly stable, kinetically trapped RNP complex that is not conducive to RNA-RNA annealing. To validate our model, we showed that alanine substitutions of the conserved acidic residues (D306, D310, E311) abrogated rotavirus replication, while charge-preserving mutations had no detrimental effect on the virus rescue in reverse genetics experiments. Our multifaceted approach provides a mechanistic basis for RNA release from high-affinity capture by NSP2, which is required for RNA annealing, chaperone dissociation, and ultimately efficient selection and packaging of a complete RV genome.

## Results

### Conserved NSP2 CTR is required for efficient RNA annealing

The C-terminal region (CTR) of NSP2 consists of a flexible linker (residues 295 − 304) that tethers an α-helix (C-terminal helix, CTH) to NSP2_core_ (i.e., residues 1 – 294, **Fig. 1a**). The CTH is ampholytic, containing highly-conserved positively-(Arg307, Lys308) and negatively-charged (Asp306, Asp310, Glu311) residues. To interrogate the role of the CTR in NSP2 function, we generated a C-terminally truncated NSP2 construct (NSP2-ΔC, residues 1 – 294) lacking the entire CTR (**fig. S1**). This NSP2-ΔC construct has been previously characterised by others (*9, 18*) (**Fig. 1**).

To visualise the CTR conformation in solution, we determined a cryo-EM 3D reconstruction of full-length NSP2 (henceforth referred to as NSP2) at a global resolution of 3.9 Å (**Fig. 1c; fig. S2 & S3**). As expected, our cryo-EM-derived model of NSP2 was highly similar to previously solved crystal structures of NSP2 (the overall RMSD between equivalent Cα atoms of the refined model presented here and PDB 1L9V is 1.124 Å) (*7, 9, 18*). Within our density map, the C-terminal helix (CTH) exhibited well-resolved density (local resolutions ranging between 3.6 – 4.0 Å (**fig. S2**)). Due to intrinsic flexibility, the linker region was poorly resolved (**fig. S3**). We confirmed that NSP2-ΔC remains octameric by determining a negative-stain EM 3D reconstruction (**fig. S1**).

Next, we investigated the role of CTR in the RNA annealing activity of NSP2 using a fluorescence cross-correlation (FCCS)-based RNA-RNA interaction assay (*17*). We chose RNA transcripts S6 and S11, representing rotavirus gene segments 6 and 11, as these have been previously shown to form stable RNA-RNA contacts in the presence of NSP2 (*17*). In brief, fluorescently-labelled rotavirus genomic transcripts (RNAs S6 and S11) were co-incubated in the absence or presence of either NSP2 or NSP2-ΔC. Ensuing intermolecular interactions were then quantitated in solution by measuring the cross-correlation function (CCF) amplitudes (**Fig. 2a**). While a zero CCF amplitude is indicative of non-interacting RNAs, increasing yields of intermolecular interactions result in proportionally higher, non-zero CCF amplitudes (*19*).

**Fig 2.**
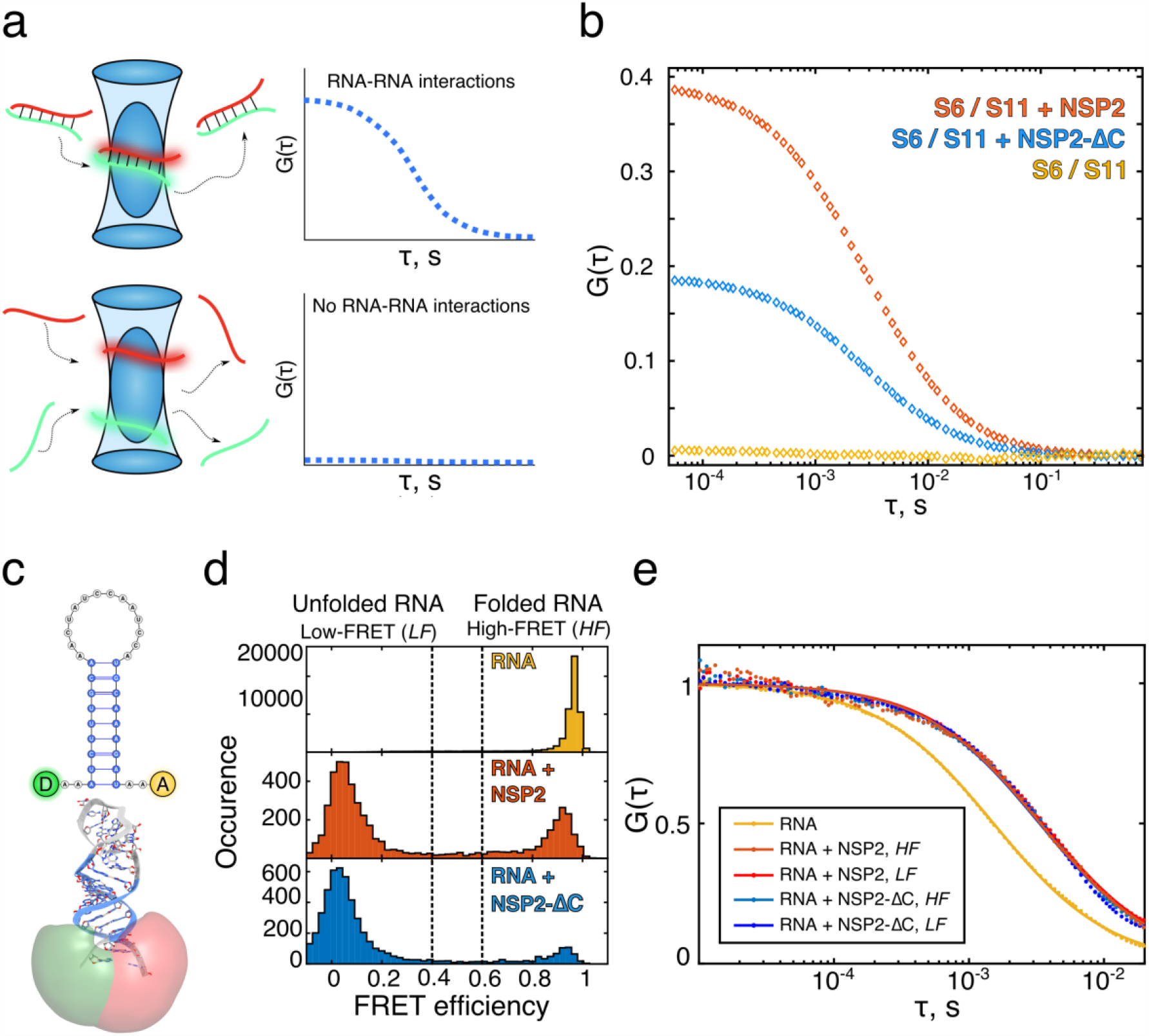
The NSP2 CTR is required for RNA-RNA matchmaking and RNA unwinding. **(a)** Single-molecule assays to probe RNA-RNA interactions between partially complementary fluorescently labelled RNAs S6 (green) and S11 (red). Upon strand annealing, differently labelled transcripts co-diffuse (shown as a duplex within the blue confocal volume). Such interactions result in a non-zero amplitude of the cross-correlation function (CCF), and thus directly report the fraction of interacting RNAs. A CCF amplitude G(τ) = 0 indicates that the two RNA molecules diffuse independently and are not interacting. **(b)** An equimolar mix of S6 and S11 RNAs were co-incubated in the absence (yellow), or presence of either NSP2 (orange) or NSP2-ΔC (blue). While the two RNAs do not interact, co-incubation with NSP2 results in a high fraction of stable S6:S11 complexes. In contrast, co-incubation of S6 and S11 with NSP2-ΔC results in 2-fold reduction of the fraction of S6:S11 complexes. **(c)** Schematics of the RNA stem-loop used for the smFRET studies of helix-unwinding activity. The FRET donor (D, green) and acceptor (A, red) dye reporters (Atto532 and Atto647N) and their calculated accessible volumes (green and red, respectively) are shown. **(d)** smFRET efficiency histograms of the RNA stem-loop alone (top, yellow), in the presence of 5 nM NSP2 (middle, red), or 5 nM NSP2-ΔC (bottom, blue). **(e)** A species-selective correlation analysis was performed on the high FRET (*HF*) and low FRET (*LF*) species of RNA stem-loops bound to NSP2 (orange) and NSP2-ΔC (blue). All FCS analyses were performed on the smFRET data shown in (d).

Co-incubation of S6 and S11 transcripts alone results in a near zero CCF amplitude indicating that they do not spontaneously interact (**Fig. 2b and fig. S3**). In contrast, addition of NSP2 to an equimolar mixture of these two RNAs produced a high CCF amplitude, indicative of intermolecular RNA duplex formation (**Fig. 2b**). This observation is in agreement with the known role of NSP2 as an RNA chaperone, facilitating the remodelling and annealing of structured RNAs (*17*).

Co-incubation of S6 and S11 in the presence of NSP2–ΔC results in a reduced CCF amplitude (**Fig. 2b**), indicating that NSP2-ΔC has a reduced RNA annealing activity relative to full-length NSP2. This is in agreement with our previous observation that NSP2-ΔC has reduced capacity to promote interactions between RV RNAs (*17*). In addition, our combined data confirms that CTR plays a role in the RNA chaperone function of NSP2 irrespective of the RNA substrates chosen.

### The NSP2 CTR reduces the RNA unwinding activity but does not directly interact with RNA

As the ability of NSP2 to unfold and remodel RNA structures is a prerequisite for its RNA annealing activity (*4*), we next investigated the role of the CTR in RNA helix destabilisation. We used single-molecule Förster Resonance Energy Transfer (smFRET) to directly compare the abilities of NSP2 and NSP2-ΔC to unwind an RNA stem-loop labelled at the 5’ and 3’ termini with donor and acceptor dyes (Atto532 and Atto647N) (**Fig. 2c**).

In the absence of either protein, the stem-loop alone adopts a folded conformation, resulting in a single, high-FRET population (*E*_FRET_ = ∼0.95) (**Fig. 2d**). Incubation with NSP2 produces two distinct FRET populations, corresponding to fully folded d(*E*_FRET_ = ∼0.95) and unfolded (*E*_FRET_ = ∼0.05) RNA states. No intermediate FRET populations (corresponding to partially-unwound stem-loop conformations) were observed, in agreement with previous observations of NSP2-mediated RNA unwinding (*4*).

We then measured the ability of NSP2-ΔC to unwind this RNA stem loop. Surprisingly, in the presence of NSP2-ΔC, the stem-loop was predominantly unfolded (*E*_FRET_ = ∼0.05) (**Fig. 2d**). Furthermore, we did not observe differences in binding of either NSP2 or NSP2-ΔC to both folded and unfolded RNA conformations (**Fig. 2e**). These data demonstrate that NSP2-ΔC has enhanced RNA unfolding activity compared to its full-length counterpart. This result is somewhat paradoxical: while NSP2-ΔC is more efficient at destabilizing RNA structure (**Fig. 2d**), it is approximately half as efficient at promoting the annealing of structured RNAs as NSP2 (**Fig. 2b**).

To deduce whether the CTR directly interacts with RNA, we used a combination of structural proteomics techniques (**Fig. 3**). We performed hydrogen-deuterium exchange-mass spectrometry (HDX-MS) experiments to map regions of NSP2 that become protected from deuterium exchange in the presence of RNA, presumably as they are involved in RNA binding and occluded from solvent when bound. We observed significant protection from exchange for peptides that predominantly mapped to ∼25 Å-deep grooves present on the surface of NSP2, indicating that this is the major RNA-binding site of NSP2 (**Fig. 3a**). Intriguingly, we did not observe any significant change in protection for peptides that spanned the CTR, indicating that the CTR does not directly interact with RNA (**Figs. 3a & 3b, fig. S5**).

**Fig 3.**
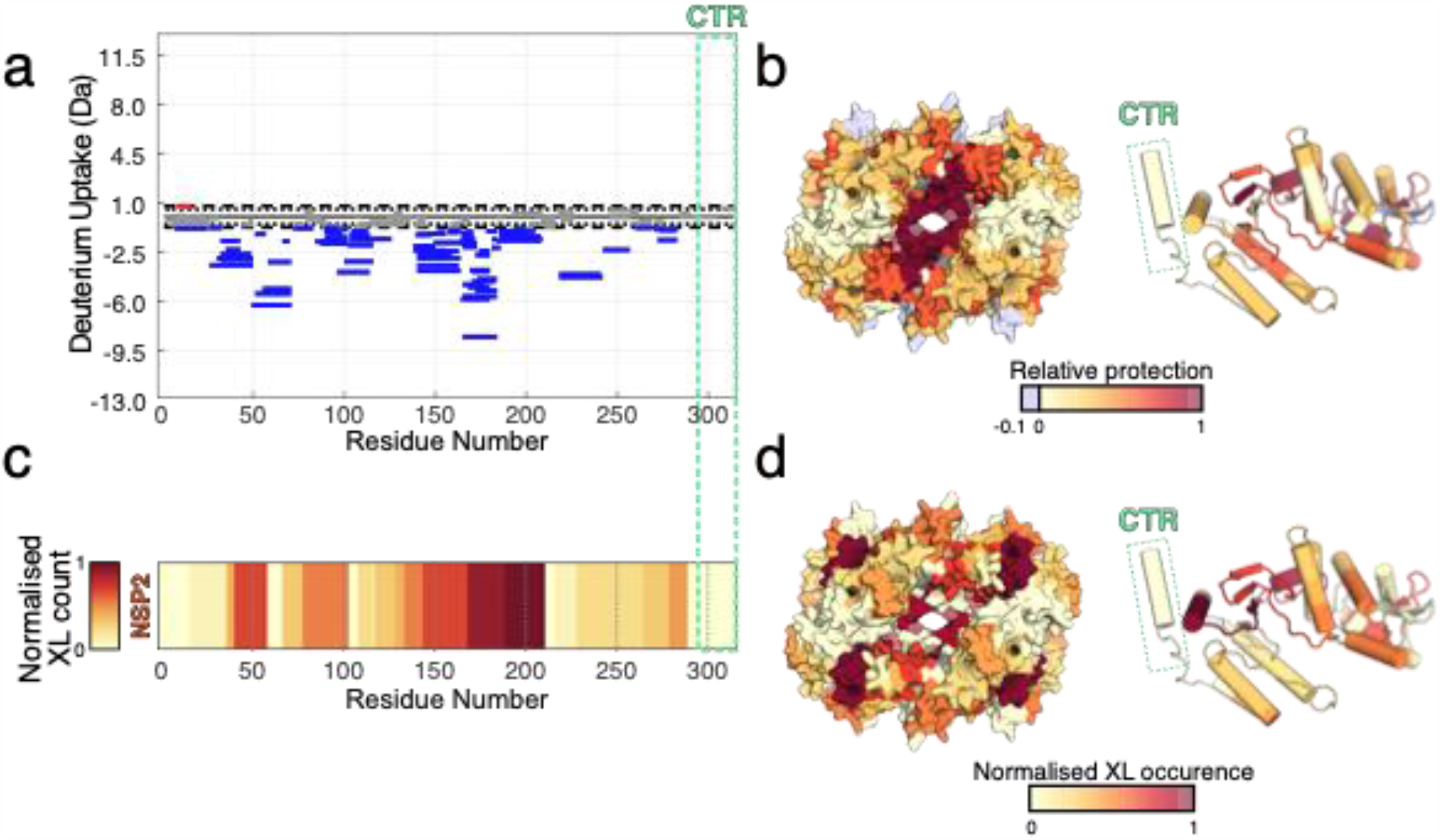
CTR does not interact with RNA. **(a)** Differences in the deuterium uptake in NSP2 (integrated over four different hydrogen-deuterium exchange timepoints), for NSP2 alone and NSP2-RNA complex. Protected and deprotected peptides are coloured blue and red, respectively. Peptides with no significant difference between conditions, determined at a 99% confidence interval (dotted line), are shown in grey. Green dashed box corresponds to the CTR, revealing no significant differences in deuterium exchange in the presence or absence of RNA. **(b)** A differential HDX map colored onto the NSP2 octamer surface (left) and monomer structure (right). Multiple regions of the NSP2 except the CTR (green box) are protected by bound RNA. **(c)** Normalised occurrence of the RNA-interacting peptides determined using UV crosslinking (identified by RBDmap). **(d)** RBDmap-identified RNA-binding peptides mapped mapped onto the surface of NSP2 octamer (left) and its monomer (right). Structures are colored according to frequency of crosslink occurrence. No RNA:peptide cross-links are mapped onto the CTR (green box).

We further corroborated the location of RNA-binding sites on NSP2 using UV-crosslinking with RBDmap (*20, 21*). Consistent with the HDX-MS data, RBDmap identified RNA-linked peptides map to these surface-exposed RNA-binding grooves (**Figs. 3c & 3d, fig. S5**). However, we again did not observe any RNA-linked peptides corresponding to the CTR.

Collectively, these results reinforce the notion that the CTR is involved in the RNA chaperone activities of NSP2. Our data indicates that although the CTR does not directly interact with RNA, it is a determinant of both the RNA unwinding and annealing activities of NSP2.

### Cryo-EM visualization of NSP2-RNA interactions

To understand the molecular basis for the RNA binding by NSP2, we determined a cryo-EM reconstruction of an NSP2 ribonucleoprotein (RNP) complex at a global resolution of 3.1 Å (**Fig. 4a, fig. S8**). While the cryo-EM density corresponding to NSP2 was well resolved, there was no density that could be attributed to the RNA in the high-resolution post-processed NSP2-RNP map (**Fig. 4a**). This is likely due to the heterogeneity and intrinsic flexibility of the NSP2-bound unstructured single-stranded RNA. Despite this, a novel feature localised to RNA binding sites identified by HDX-MS and RBDmap (**Fig. 3**) was present in 5 Å low-pass filtered (LPF) maps (**Fig. 4b**). Notably, such a density feature was not present in 5 Å-LPF NSP2 apoprotein maps (**Fig. 4c**). We attribute this density to NSP2-bound RNA.

**Fig 4.**
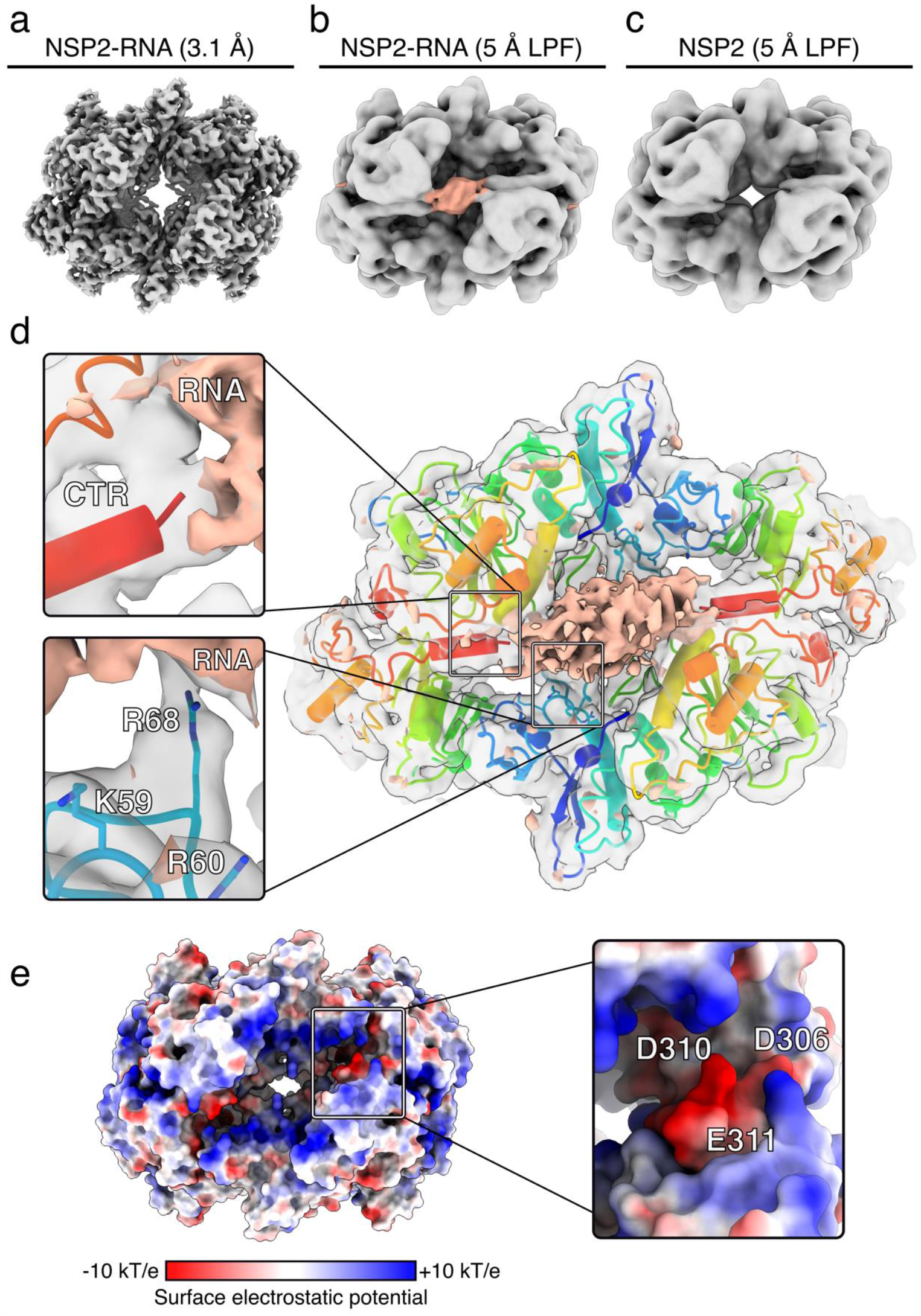
Cryo-EM structure of the NSP2-RNP complex. **(a)** 3.1 Å-resolution reconstruction of the NSP2-RNP complex. (**b-c)** Cryo-EM maps of NSP2-RNA (**b**) and NSP2 apoprotein (**c**) low-pass filtered (LPF) to 5 Å. A novel cryo-EM density feature (peach) is attributed to bound RNA in the LPF RNP map. Both maps are reconstructed with D4 symmetry. **(d)** Direct visualization of interactions between NSP2 and RNA using C4 symmetry expansion and focused classification. The positive difference density map corresponding to RNA (peach) is overlaid onto the unsharpened NSP2-RNP complex map determined through symmetry expansion and focused classification (grey, transparent density) and atomic model of NSP2. Inset: zoom in of CTR positioned relative to RNA density (top) and RNA-interacting residues (bottom). (**e**) The surface electrostatic potential analysis of NSP2. Inset: zoom-in of the CTR, with residues within the acidic patch (D306, D310, E311) annotated.

To improve visualization of the RNA density, we performed focused classification using C4 symmetry-expanded data with a mask applied to a single RNA-binding face of NSP2 (*22*–*24*). The resulting 3D reconstructions readily classified into four dominant populations, three of which had poor RNA occupancy (each class with 26% of the input particles), while a single 3D class average (22% of input particles) exhibited improved RNA density (**fig. S2**). Due to the reasons outlined above, the diffuse nature of this density prevented us from modelling the ssRNA into the structure. However, we were able to visualise residue-specific NSP2-RNA contacts (**Fig. 4d**).

We built an atomic model of NSP2 into the sharpened map and then computed a difference map between NSP2 and the RNA-occupied, focused map in order to visualise the NSP2-RNA contacts. Significant positive density was localised in the basic groove of NSP2 (**Fig. 4**), consistent with the binding site identified through HDX and RBDmap (**Fig. 3**). We observed interactions between positively-charged residues, most notably R68 (**Fig. 4d**). Adjacent to this contact are K58, K59, and R60, of which K59 and R60 are directly oriented towards the RNA density (**Fig. 4d**, inset). The importance of these residues for RNA capture by NSP2 is strongly supported by previous biochemical studies that identified a number of solvent-exposed lysine and arginine residues (K37, K38, K58, K59, R60, R68) that span the periphery of the NSP2 octamer (**fig. S6**) and contribute to RNA binding (*25*).

Furthermore, the identified residues are localised to an unstructured loop within the RNA-binding groove, allowing promiscuous and flexible accommodation of alternative RNA structures with near-identical affinities by NSP2, consistent with previous reports (*4, 26*). Together with our HDX and RBDmap results, our cryo-EM reconstruction reveals a number of electrostatic contacts that provide a plausible molecular basis for non-specific NSP2-RNA interactions (**Fig. 4d**, inset). In addition, our EM reconstruction has revealed a number of other residues (R240, K286, F290) that likely contribute to RNA binding, also identified by HDX and RBDmap (**figs. S5 & S6**). These residues may also participate in non-specific RNA contacts via electrostatic interactions, hydrogen bonding and π-π stacking, consistent with a significant non-electrostatic contribution to the overall free energy of RNA binding to NSP2.

### Conserved acidic patch within the CTR promotes RNA dissociation

Within the cryo-EM density map, the CTRs are poised below the RNA, while making limited contacts with the observed RNA density (**Fig. 4d**). This suggests that rather than modulating the RNA binding affinity, the CTR may play a role in promoting RNA dissociation from NSP2. To investigate this, we performed binding kinetics measurements using surface plasmon resonance (SPR) (**Figs. 5a & b**). Association rate constants (*K*_on_) remain largely consistent across a range of concentrations of both NSP2 and NSP2-ΔC (NSP2-ΔC binds 1.5 ± 0.4-fold faster than NSP2 (**table S1**)). However, NSP2-ΔC exhibited 3.2 ± 0.3-fold slower dissociation than NSP2, suggesting a role for CTR in the displacement of bound RNA (**Fig. 5a, table S1**).

**Fig 5.**
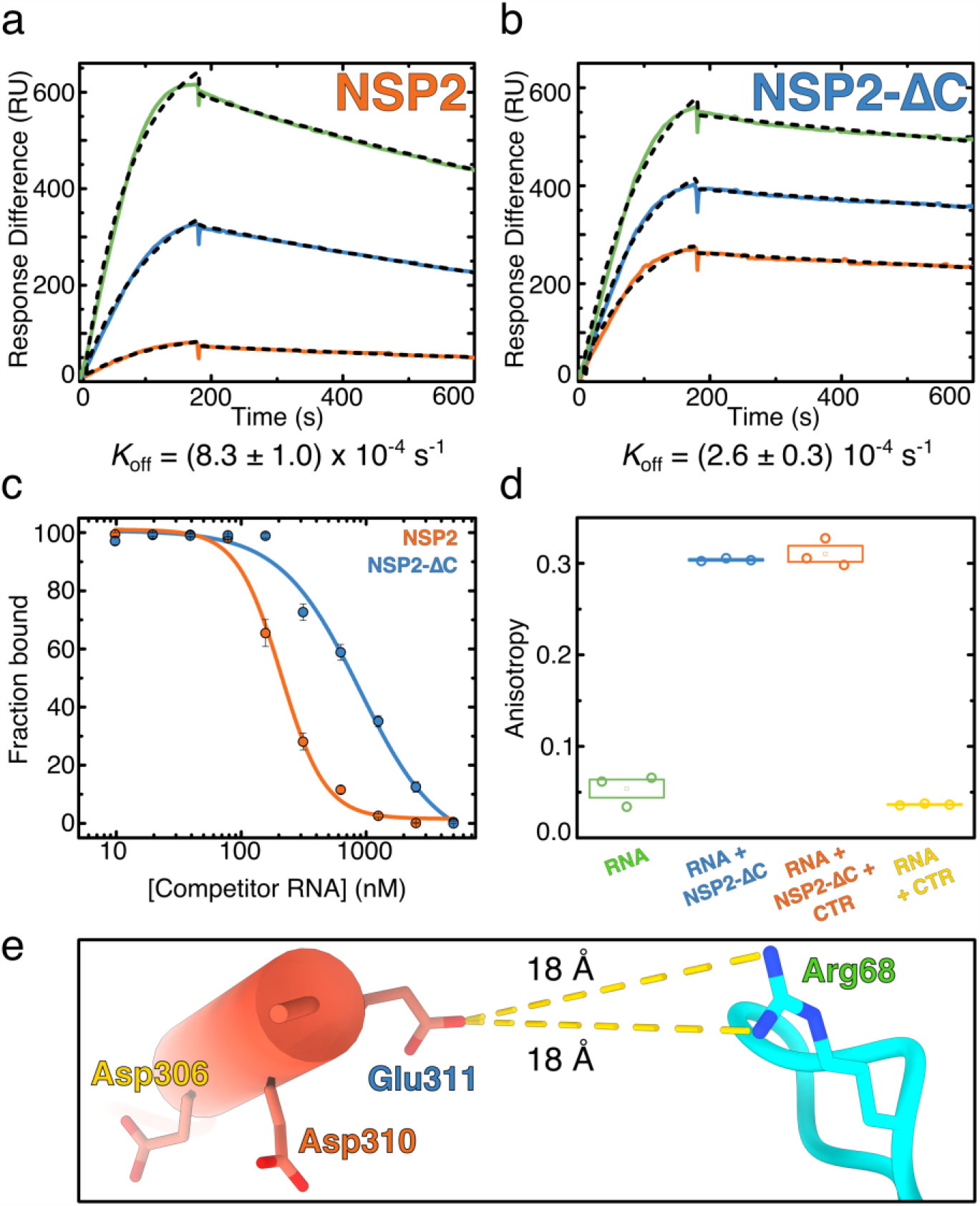
The CTR promotes RNA dissociation non-competitively. **(a-b)** SPR sensograms of NSP2 (**a**) and NSP2-ΔC (**b**) binding to RNA. Although NSP2-ΔC binds RNA with ∼6-fold higher affinity, this is due to a modest (1.5-fold) increase in *K*_on_, and a larger (3.2-fold) decrease in *K*_off_. **(c)** RNA competition assay with the CTR peptide. The fractional binding of fluorescently labelled RNA was determined by fluorescence anisotropy. Labelled RNA (10 nM) fully bound to NSP2 (orange) or NSP2-ΔC (blue) was titrated with unlabelled RNA of identical sequence to compete for NSP2 binding against labelled RNA. The IC_50_ values for NSP2 and NSP2-ΔC are 208 ± 11 nM and 890 ± 160 nM, respectively. The NSP2-RNA complex undergoes strand exchange more readily than the NSP2-ΔC:RNA complex. **(d)** RNA binding by NSP2 in the presence of the CTR peptide. CTR peptide (10 µM) was added to preformed NSP2-ΔC:RNA complexes. The CTR peptide does not compete with RNA for binding to NSP2-ΔC. (**e**) Estimated distances between acidic residues within CTR and the R68 that interacts with RNA. Note the nearest side chain (E311) which is 18Å away from R68.

Close examination of our cryo-EM-derived model revealed an acidic patch within the CTR, (**Figs. 4e & f**). This is in contrast to other clusters of surface-exposed acidic residues on NSP2 that show a low degree of conservation (**fig. S10**). The acidic patch of the CTR is presented directly underneath the density attributed to bound RNA (**Fig. 4d**), potentially promoting RNA displacement from the NSP2. Such displacement could be achieved either via direct competition with RNA-binding residues or by providing a negatively-charged environment that accelerates RNA dissociation from NSP2 through charge repulsion.

Therefore, to further investigate RNA displacement from NSP2, we used RNA competition assays (**Figs. 5c & d**). We performed titrations of unlabelled RNA into preformed RNP complexes containing fluorescently-labelled RNA to understand the differences in RNA exchange and chaperone recycling between NSP2 and NSP2-ΔC. Using fluorescence anisotropy, we estimated the degree of competition as the concentration of competitor RNA required to displace 50% of pre-bound RNA from either NSP2 or NSP2-ΔC complexes (IC_50_). We determined IC_50_ values of 208 ± 11 nM and 890 ± 160 nM for NSP2 and NSP2-ΔC, respectively, confirming that NSP2–ΔC undergoes ∼4-fold reduced RNA exchange, consistent with its ∼3-fold slower rate of dissociation from RNA (**Figs. 5a & b**).

We then investigated whether the CTR promotes RNA dissociation from NSP2 through directly competing with RNA for binding to basic, RNA-binding residues on the NSP2_core_. To achieve this, we measured RNA binding by NSP2-ΔC in the presence of saturating amounts of a synthetic peptide matching the sequence of the CTR. No dissociation of RNA from the NSP2-ΔC was observed in the presence of 20-fold molar excess of the CTR peptide over NSP2-ΔC (**Fig. 5d**). Furthermore, no RNA binding was observed upon incubation with 10 µM CTR peptide (i.e. 400-fold excess), indicating that the CTR does not bind RNA. This suggests that, while the CTR is required for RNA displacement from NSP2, this does not occur through direct competition.

We analysed our atomic model of NSP2 to evaluate the distances between acidic residues within the CTR and the basic, RNA-binding residues localised to flexible loops within the RNA-binding grooves (**Fig. 5e & fig. S6**). The distances (∼10 – 30 Å) between acidic residues within the CTR and the RNA-interacting residues are incongruent with a direct competition model. While R68 was demonstrated to directly interact with RNA (**Fig. 4d**), it is 18 Å away from acidic residues within the CTR. This further demonstrates that, while CTR promotes dissociation of RNA from NSP2, it does not do so through direct competition for NSP2_core_ binding (**Fig. 5e**). Collectively, our data suggest that conserved acidic patches within the CTR promote dissociation of bound RNA from NSP2 via charge repulsion.

Finally, to validate our findings *in vivo*, we employed a reverse genetics approach to rescue recombinant rotaviruses with point mutations within the CTR. We assessed the effects of amino acid substitutions within the CTR on viral replication by attempting recombinant virus rescue, as described in Materials and Methods. All attempts to rescue a triple alanine mutant D306A/D310A/E311A were unsuccessful, suggesting these mutations completely abrogate virus replication. Remarkably, a triple mutant containing charge-preserving mutations D306E/D310E/E311D was successfully rescued with the same efficiencies as wild-type virus (**Fig. S7**). This directly demonstrates the essential role of the CTR acidic patch in rotavirus replication.

## Discussion

Long RNAs adopt an ensemble of diverse stable structures that limit spontaneous RNA-RNA interactions through the sequestration of sequences required for intermolecular base pairing (*17, 27*–*30*). This necessitates the action of RNA chaperone proteins to bind and refold RNA structures in order to promote RNA annealing between complementary sequences (*31*–*33*).

In order to function as an RNA chaperone, NSP2 must capture, unwind, anneal and release complementary RNA sequences (*17, 34, 35*). Previous structural studies have provided static snapshots of crystallographically-averaged NSP2-RNA complexes (*7, 18, 25*). However, due to the highly dynamic nature of the protein-RNA interactions required for its RNA chaperone activity, they have only revealed limited insights into the molecular mechanisms of NSP2. Our recent work (13, 17, 18) indicates that for NSP2-RNP complexes such heterogeneity arises from poorly defined protein-RNA stoichiometries and the ability of bound RNA to adopt multiple configurations and orientations. To overcome these challenges, here we used a combination of single molecule fluorescence, cryo-EM, structural proteomics and biophysical assays to decipher the mechanism of NSP2 chaperone function.

Previous work suggests that the C-terminal region of NSP2 is essential for rotavirus replication (*15*). Using single-molecule fluorescence techniques, here we have directly shown that the CTR of NSP2 is important for promoting RNA-RNA interactions. However, we only identified interactions between RNA and basic residues located in flexible loops within the RNA binding groove of NSP2, but not the CTR. Remarkably, similar RNA recognition mechanisms have been reported in other RNA chaperones including *E. coli* StpA and HIV-1 NC (*37, 38*). Collectively, these results highlight the role of the CTR in NSP2 RNA chaperone activity but not RNA binding.

### Mechanism of the CTR-assisted RNA displacement and its role in RNA matchmaking

FCS analysis of high- and low-FRET RNA species points out that full length NSP2 binds to both the unfolded and folded RNA conformations, priming RNAs for efficient RNA annealing (**Fig. 3e & fig. S4**). Single-molecule fluorescence and binding kinetics experiments indicate that removal of the CTR does not perturb RNA binding but slows RNA release (∼3.2-fold increase in k_off_). Moreover, CTR removal results in a ∼2.4-fold increase in the RNA-unwinding activity of NSP2-ΔC, as well as a ∼2-fold decrease in its RNA annealing activity. Additionally, smFRET data reveal that binding to NSP2-ΔC energetically favours low-FRET (unfolded) RNA conformations, resulting in remodelling of structured RNAs (**Fig. 3d**). The resulting increased stability of NSP2-ΔC-RNA complexes precludes efficient RNA annealing, yielding kinetically trapped RNP complex intermediates.

We propose a model whereby rotavirus RNA chaperone NSP2 binds to RNA with high affinity, resulting in RNA structure destabilisation (**Fig. 6a**). By binding to multiple RNAs concurrently via surface-exposed grooves (**Fig. 6a**, cyan) (*4, 7, 17*), it acts as a matchmaker of complementary sequences, promoting intermolecular RNA-RNA interactions. Conserved acidic patches within the ampholytic CTR (**Fig. 6a**, red) accelerate RNA displacement from NSP2 via charge repulsion, thus enabling RNA chaperone recycling and duplex release. Removal of the ampholytic CTRs in the NSP2 variants derived from two distinct rotavirus viruses (strains SA11 and RF) has a similar outcome on RNA chaperone activity *in vitro* (**fig. S3**), suggesting a conserved role of the CTR in NSP2 function. This model is further supported by our observation that removal of the unstructured region downstream of the ampholytic CTR (**Fig. 1a**) does not alter the RNA unwinding activity of NSP2 (**fig. S4**). Indeed, this partial truncation has been previously shown to support viral replication (*9*). Excitingly, we have exploited recent technical advances in reverse genetics of rotaviruses to directly demonstrate the pivotal role of these conserved acidic residues in rotavirus replication (**fig. S7**).

**Fig 6.**
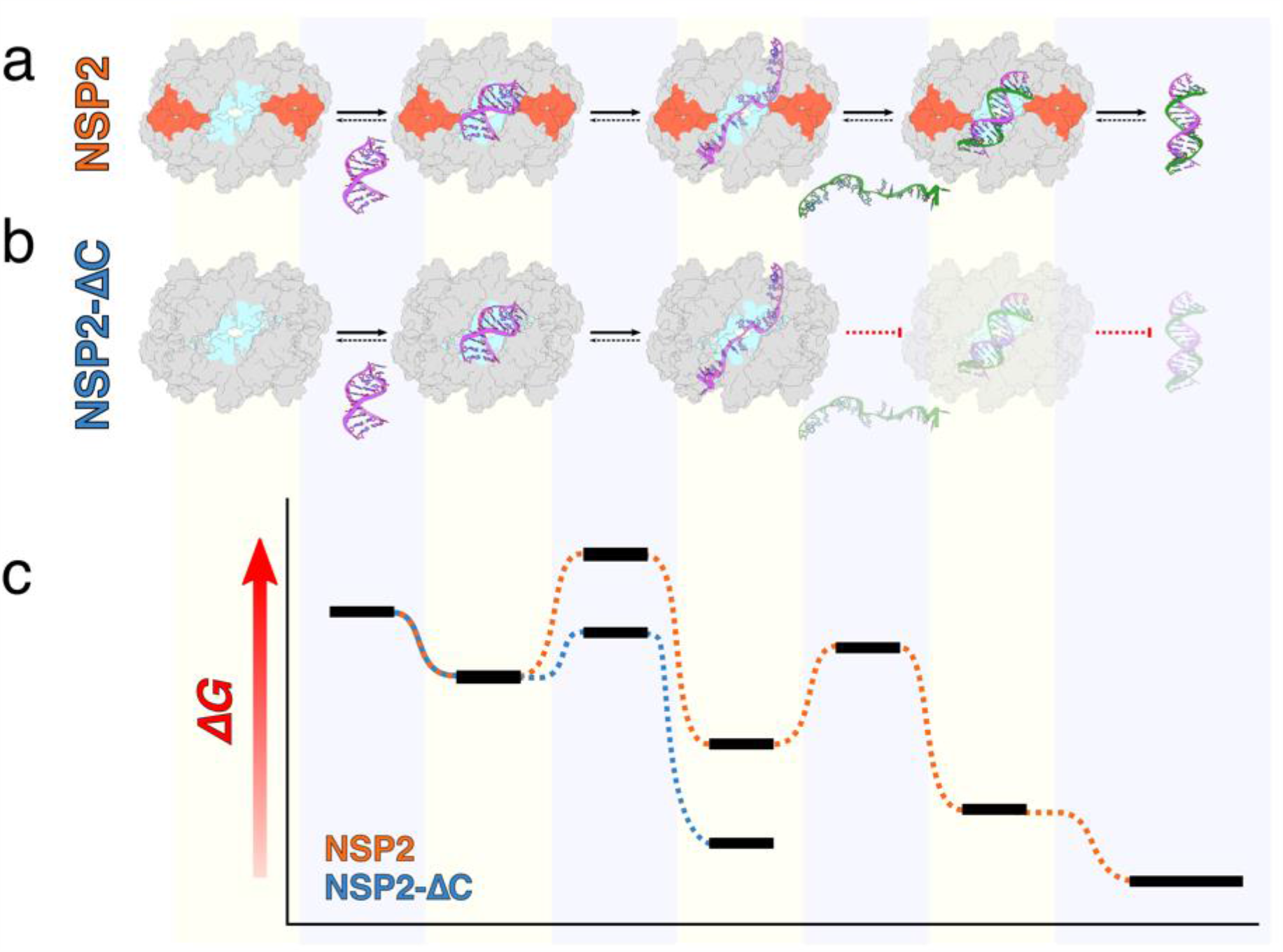
Proposed mechanism of CTR-accelerated RNA dissociation and its requirement for efficient NSP2-mediated RNA-RNA interactions. **(a)** NSP2 captures RNA (purple) via a positively-charged groove (cyan) and promotes RNA unwinding. Binding of a second, complementary RNA strand (green) by NSP2 allows efficient annealing and the proximity to the CTR (burnt orange) promotes dissociation of dsRNA from the NSP2. **(b)** In contrast, NSP2-ΔC captures and unwinds RNA, forming a highly stable intermediate. The stability of the intermediate state makes displacement of the bound RNA by a complementary RNA segment via annealing thermodynamically unfavourable. **(c)** A free energy diagram of NSP2 (orange) and NSP2-ΔC (blue)-mediated RNA annealing. Horizontal black bars correspond to the free energy levels of different RNA states corresponding to the above schematic representations in (a) & (b).

The described principle of CTR-assisted RNA dissociation from NSP2 is strikingly similar to that of the bacterial RNA chaperone protein Hfq (*39*–*41*). Hfq possesses an unstructured C-terminal domain (CTD) with an acidic tip that drives RNA displacement from the Hfq_core_ (*42*). Unlike the Hfq CTD, we do not observe competition between CTR and RNA for NSP2_core_ binding (*43, 44*). Nevertheless, the NSP2 CTR modulates the kinetics and thermodynamics of NSP2-RNP complex formation to accomplish RNA chaperone recycling. We propose that this may represent a conserved mechanistic feature of multimeric RNA chaperones that capture RNA with high affinity and require auto-regulation to assist RNA dissociation in order to promote efficient matchmaking.

## Materials and Methods

### Protein and RNA production

NSP2 and NSP2-ΔC (RVA strains SA11 and RF) were expressed and purified as described previously (*17*). RNAs used in this study are listed in **table S2**. RNA sequences S5, S6 and S11 were produced and labelled using previously-described *in vitro* transcription and labelling protocols (*17*). Unstructured 20mers (labelled and unlabelled), unstructured 10mer (labelled) and biotinylated unstructured 10mer RNAs were purchased from Integrated DNA technologies (IDT). Dual-labelled stem loop RNA was purchased from IBA Life Sciences.

### Negative stain electron microscopy and data processing

For negative-stain grid preparation, 4 µl of sample (at various concentrations ranging from 100 – 500 nM) was incubated on glow-discharged (using PELCO easiGlow) carbon-coated Formvar 300-mesh Cu grid (Agar scientific) for 90 seconds prior to blotting, and stained twice with 20 µl 2% uranyl acetate (first stain immediately blotted, the second stain incubated for 20 seconds prior to blotting) and allowed to dry. Micrographs were collected on a FEI Tecnai 12 transmission electron microscope operated at 120 kV and equipped with a Gatan UltraScan 4000 CCD camera operated at a nominal magnification of 30,000 x (giving a 3.74 Å / pixel sampling on the object level). From 23 micrograph images taken with a nominal defocus of − 3 µm, 14,740 particles were picked using template-based autopicking within Relion 3. Multiple rounds of 2D and 3D classification resulted in the selection of a subset of 2,864 particles. These particles were used to determine a ∼22 Å resolution NSP2-ΔC reconstruction with D4 symmetry applied.

### Cryogenic electron microscopy (cryo-EM) and data processing

Cryo-EM was performed exclusively with Quantifoil R.1.2/1.3 holey carbon grids (i.e. regular ∼1.2 µm circular holes with a regular spacing of ∼1.3 µm), purchased from Quantifoil. All grids were glow discharged in air using GloQube glow discharge system (Quorum) immediately prior to use. All grids were prepared using a Vitribot IV (FEI) at 100% humidity and 4°C, with a blotting time of 6 seconds and a nominal blotting force of 6. Samples were flash-frozen in liquid nitrogen (LN_2_)-cooled liquid ethane and immediately transferred to storage dewars under LN_2_.

Vitrified samples were imaged at low temperature in-house (Astbury Biostructure Laboratory, University of Leeds), using Thermo Fisher Titan Krios microscopes equipped with either a Falcon III (NSP2 apoprotein) or a Gatan K2 (NSP2-RNP) detector. Data was collected with an acceleration voltage of 300 kV and a nominal magnification of 75,000x, resulting in pixel sizes of 1.065 Å (Falcon III) or 1.07 Å (K2). Data collection parameters are described in **table S3**.

Image processing was carried out using the Relion 3 pipeline (*45*). Movie drift-correction was performed using MOTIONCOR2 (*46*), and the contrast transfer function of each movie was determined using gCTF (*47*). Initial particle autopicking of a subset of 5 – 10 randomly chosen micrographs was performed with the Laplacian-of-Gaussian (LoG) tool within the Autopicking module of Relion3. Particles were extracted and subjected to initial 2D classification in order to identify particles and assess autopicking success. Following this, the entire dataset was picked using LoG methods, extracted using 256 pixel box size and binned four times (effective box size 64 pixels) and subjected to 2D classification with fast subsets in order to remove false-positive particles that had been erroneously picked. Next, a more rigorous 2D classification was performed (without fast subsets). Particles originating from 2D classes with secondary structural features were selected and used to generate an initial model. Following multiple rounds of 3D and 2D classification, suitable particles were selected for 3D auto-refinement and various symmetry parameters were applied. Following refinement, per-particle CTF and Bayesian Polishing were performed in Relion 3, and ‘shiny’ particles were re-refined. Post-processing was performed with a soft mask of 15 pixels and the B-factor estimated automatically in Relion 3.

After particle polishing, the NSP2 apoprotein and the NSP2-RNP complex were subjected to further 3D classification into three classes without particle orientations. This yielded three similarly-sized subsets of near-identical particles for the NSP2 apoprotein whose resolution did not improve upon the original, larger dataset following 3D auto-refinement and post-processing. For the NSP2-RNP complex, this gave a single class with 86% of particles, and two other classes with 6% and 8% of the particles. The class with 6% of input particles had well-defined protein and RNA densities and was used for 3D auto-refinement and post-processing. This improved the map resolution from 3.5 Å to 3.4 Å with C4 symmetry. A D4 symmetry reconstruction further increased the map resolution from 3.5 Å to 3.1 Å.

For the NSP2 RNP complex, symmetry expansion was performed on a subset of the 635,599 particles used for a C4 symmetry reconstruction using the *relion_particle_symmetry_expand* command, generating four symmetry-related orientations for each particle. A mask covering a single basic groove-face of NSP2 was made using the volume eraser tool in UCSF ChimeraX (*48*) and Relion 3, with a soft edge of 15 pixels (**fig. S8**). The symmetry-expanded dataset was then subjected to focussed classification into 10 classes using this mask without particle orientations. Suitable classes (four classes containing >99% of input particles) were selected, and manually examined for putative RNA density. The subset of particles with the strongest RNA density feature were reconstructed without a mask, and subjected to masking (with the mask corresponding to the entire NSP2 octamer rather than a single face) and post-processing as described for reconstructions with D4 symmetry imposed. Sharpened asymmetric and D4 symmetry maps were aligned using the Fit-In-Map tool within UCSF Chimera and had a correlation of 0.9663.

### Atomic model building

A previous atomic model of NSP2 (PDB 1L9V) was fit into the cryo-EM densities using ChimeraX (*48*), and subjected to automated flexible fitting and refinement using Namdinator (*49*). The Namdinator model was used for multiple iterative rounds of manual adjustment in Coot (*50*) and real-space refinement in Phenix (*51*). Models for NSP2 apoprotein and NSP2-RNP were validated using MolProbity (*52*) as implemented in Phenix.

### Protein Sequence Conservation Analysis

Full-length NSP2-coding sequences of group A rotaviruses were obtained from GenBank. Sequences of avian strains and of rearranged RNA segments of mammalian strains were excluded from the analysis. Protein sequence conservation and multiple sequence alignment (MSA) was performed using the online ConSurf (*53*) server. Output from the ConSurf MSA was used to generate a sequence logo using the WebLogo server (*54*). Maps and models were visualized using ChimeraX (*48*) and the electrostatic surfaces were determined using the APBS plugin (*55*).

### Single-molecule (sm)FRET measurements

SmFRET measurements of freely diffusing dual-labeled RNA stem-loops in the presence and absence of NSP2 were performed on a home-built confocal microscope as described previously (*4*). Briefly, the samples were excited using pulsed interleaved excitation (*56*) at wavelengths of 532 and 640 nm (PicoTA, Toptica and LDH-D-C-640, PicoQuant) with typical laser powers of 100 µW as measured before the 60x water immersion objective (Plan Apo IR 60x/ 1.27 WI Nikon, Düsseldorf, Germany). The fluorescence signal was split between the green and red detection channels using a DualLine Z532/635 beamsplitter (AHF) and the emission spectra filtered using a Brightline 582/75 filters (Semrock) for green detection and HQ700/75 and ET700/75 filters (Chroma) for red detection. Measurements were performed in eight-well chamber slides (Nunc Lab-Tek, VWR) in a buffer composed of 1/3 PBS (45 mM NaCl, 3 mM phosphate, 1 mM KCl), 1 mM Trolox to reduce photobleaching (*57*) and 0.01% (v/v) Tween20 to prevent sticking of the sample to the glass surface. The dual-labeled RNA stem-loop was diluted to 25 pM and incubated with 5 nM NSP2 (either full length or the ΔC mutant). Data were analyzed with the open-source software package PAM (*58*) using the same burst search parameters and correction factors as described in (*4*). To determine species-selective fluorescence correlation functions, we defined two sub-populations based on the FRET efficiency *E*: the low-FRET population with *E* < 0.4 and the high-FRET population with *E* > 0.6. For each burst, the correlation function for acceptor photons after acceptor excitation was calculated including photons within a time window of 20 ms.

### Surface plasmon resonance (SPR)

A Biacore 3000 was used to analyse the binding kinetics of NSP2 and NSP2-ΔC to 5’biotinylated-10mer RNA (**table S2**). All experiments were performed in SPR buffer (150 mM NaCl, 25 mM HEPES, pH 7.5, 0.1 % Tween-20). RNAs were immobilized on an SA sensor chip (GE Healthcare) with an analyte R_*max*_ of ∼20 resonance units (RU). Analyte measurements were performed at 25°C and a flow rate of 40 µL/min. The chip surface was regenerated between protein injections with a 40 µl 0.05% SDS injection. Data were analysed using BIAevaluation 3.1 software (GE Healthcare). The kinetic parameters were derived assuming a binding stoichiometry of 1: 1.

### RNA competition assay

250 nM NSP2 and NSP2-ΔC (RF) were pre-incubated with 10 nM 20mer AlexaFluor488-labelled RNA (**table S2**) in binding buffer (50 mM NaCl, 25 mM HEPES pH 7.5). Fluorescence anisotropy measurements were performed in the presence of various concentrations of unlabelled 20mer RNA in low-volume Greiner 384-well plates. Data were recorded at 25°C in a PHERAstar Plus multi-detection plate reader (BMG Labtech) equipped with a fluorescence polarization optical module (*λ*_ex_ = 485 nm; *λ*_em_ = 520 nm). The data were normalised and binding curves were fitted in Origin 9.0 using a Hill binding curve resulting in R^2^ values of 0.997 and 0.991 for NSP2 and NSP2-ΔC respectively.

### CTR peptide competition assay

In order to maximise any potential competition between the CTR peptide and RNA, assays were performed under conditions that favoured dissociation of NSP2-ΔC from RNA. The binding assay was performed in PBS buffer (150 mM NaCl, 10 mM potassium phosphate, 3 mM potassium chloride). 25 nM AlexaFluor488-labelled RNA was incubated with 20-fold excess NSP2-ΔC (500 nM, RF strain). After 30 minutes at room temperature (∼25°C), the CTR peptide was added in 20-fold excess of NSP2-ΔC (i.e. 10 µM). To investigate direct CTR-RNA interactions, 25 nM 10mer RNA was also co-incubated with 10 µM CTR peptide. Fluorescence anisotropy measurements of RNA alone, RNA-NSP2-ΔC, RNA:NSP2-ΔC:CTR and RNA:CTR were performed in triplicate as described above for RNA competition assays.

### Hydrogen-deuterium exchange mass spectrometry (HDX-MS)

An automated HDX robot (LEAP Technologies, Ft Lauderdale, FL, USA) coupled to an Acquity M-Class LC and HDX manager (Waters, UK) was used for all HDX-MS experiments. Differential HDX-MS of NSP2 was performed using NSP2 (10 µM) or pre-incubated NSP2-RNP complexes (10 µM + 2 µM 20mer RNA, **table S2**). 30 µl of protein-containing solution was added to 135 μL deuterated buffer (10 mM potassium phosphate buffer pD 8.0, 82% D_2_O) and incubated at 4 °C for 0.5, 2, 30 or 120 min. After labelling, HDX was quenched by adding 100 μL of quench buffer (10 mM potassium phosphate, 2 M Gdn-HCl, pH 2.2) to 50 μL of the labelling reaction. 50 μL of the quenched sample was passed through immobilised pepsin and aspergillopepsin columns (Affipro, Mratín, Czech Republic) connected in series (20 °C) and the peptides were trapped on a VanGuard Pre-column [Acquity UPLC BEH C18 (1.7 μm, 2.1 mm × 5 mm, Waters, UK)] for 3 min. The peptides were separated using a C18 column (75 μm × 150 mm, Waters, UK) by gradient elution of 0–40% (v/v) acetonitrile (0.1% v/v formic acid) in H_2_O (0.3% v/v formic acid) over 7 min at 40 μL min^−1^. Peptides were detected using a Synapt G2Si mass spectrometer (Waters, UK). The mass spectrometer was operated in HDMS^E^ mode with the dynamic range extension enabled (data independent analysis (DIA) coupled with IMS separation) were used to separate peptides prior to CID fragmentation in the transfer cell. CID data were used for peptide identification and uptake quantification was performed at the peptide level (as CID results in deuterium scrambling). Data were analysed using PLGS (v3.0.2) and DynamX (*59*) (v3.0.0) software (Waters, UK). Restrictions for peptides in DynamX were as follows: minimum intensity = 1000, minimum products per amino acid = 0.3, max sequence length = 25, max ppm error = 5, file threshold = 3. The software Deuteros (*60*) was used to identify peptides with statistically significant increases/decreases in deuterium uptake (applying a 99 % confidence interval) and to prepare Woods plots.

### UV-crosslinking-mass spectrometry with RBDmap

10 µM NSP2 was incubated with 5’-A_25_-S11 RNA in a final volume of 100 µl. NSP2-RNP complexes were incubated at room temperature for 30 minutes and applied to a single well of a 24-well plate. This 24-well plate was placed on an aluminium block cooled to 4°C within a plastic container of ice and subjected to 6 rounds of UV irradiation (254 nm, 0.83 J cm^-2^ per round) in a UVP CL-1000 Ultraviolet Crosslinker (Scientifix). Crosslinked RNP complexes were digested by LysC (NEB, #P8109S) (500 ng per crosslinked RNP sample) overnight at room temperature. Enrichment and identification of cross-linked peptides were performed using the in vitro adaptation of the RBDmap protocol, as described in (*20*). Data analysis was performed using the CrissCrosslinker R script, as described in (*20*).

### Circular dichroism (CD)

CD experiments were performed in a Chirascan plus spectrometer (Applied Photophysics). Samples were prepared by dialyzing protein solutions against 10 mM phosphate buffer pH 7.4, 50 mM sodium fluoride. Spectra were recorded over a wavelength range of 190–260 nm with a bandwidth of 1 nm, step size of 1 nm and a path length of 1 mm. An average of three scans were used for the final spectra.

### NSP2 (RF) threading and sequence alignment

Alignment of NSP2(SA11) and NSP2(RF) sequences was performed using T-Coffee (*61*). Threading of the NSP2(RF) sequence (based on the SA11 structure, PDB 1L9V) was performed using I-TASSER (*62*).

### Cells and viruses

MA104 (embryonic African green monkey kidney cells, ATCC® CRL-2378) were cultured in Dulbecco’s Modified Eagle’s Medium (DMEM) (Life Technologies) supplemented with 10% Fetal Bovine Serum (FBS) (Life Technologies). BHK-T7 cells (Baby hamster kidney stably expressing T7 RNA polymerase) were cultured in Glasgow medium supplemented with 5% FBS, 10% Tryptose Phosphate Broth (TPB, Sigma-Aldrich), 2% Non-Essential Amino Acid (NEAA, Sigma) and 1% Glutamine. Recombinant simian RV strain SA11 NSP2 mutants were rescued using cDNA clones encoding the wild-type SA11 (G3P[2]) virus with modifications, as previously described. Briefly, plasmids pT_7_-VP1-SA11, pT_7_-VP2-SA11, pT_7_-VP3-SA11, pT_7_-VP4-SA11, pT_7_-VP6-SA11, pT_7_-VP7-SA11, pT_7_-NSP1-SA11, pT_7_-NSP2-SA11, pT_7_-NSP3-SA11, pT_7_-NSP4-SA11, and pT_7_-NSP5-SA11 (*11*) were used for recombinant virus rescue in reverse genetics experiments. pT_7_-NSP2 plasmid variants (NSP2-EED, NSP2-AAA, and NSP2-6xHis) carrying mutations in gs8 were generated using Q5 site-directed mutagenesis (NEB), using mutagenesis primers (Table S2). To rescue recombinant RV strain SA11, monolayers of BHK-T7 cells (4 × 10^5^) cultured in 12-well plates were co-transfected using 2.5 μL of TransIT-LT1 transfection reagent (Mirus) per microgram of DNA plasmid. Each mixture comprised 0.8 μg of SA11 rescue plasmids: pT_7_-VP1, pT_7_-VP2, pT_7_-VP3, pT_7_-VP4, pT_7_-VP6, pT_7_-VP7, pT_7_-NSP1, pT_7_-NSP3, pT_7_-NSP4, and 2.4 μg of pT_7_-NSP2 and pT_7_-NSP5 (*11, 63*). Additional 0.8 μg of pcDNA3-NSP2 and 0.8 μg of pcDNA3-NSP5, encoding NSP2 and NSP5 proteins, were also co-transfected to increase the virus rescue efficiencies. At 24 h post-transfection, MA104 cells (5 × 10^4^ cells) were added to the transfected cells and co-cultured for 72 hours in FBS-free medium supplemented with trypsin (0.5 μg/mL, Sigma Aldrich). After incubation, transfected cells were lysed by repeated freeze-thawing and 0.2 ml of the lysate was transferred to a fresh MA104 cell monolayer. After adsorption at 37°C for 1 hour, followed by a 5 min wash with PBS, cells were further cultured for 4 days in FBS-free DMEM supplemented with 0.5 μg/mL trypsin until a clear cytopathic effect (CPE) was visible. For AAA mutant, where CPE was not observed after 4 days of incubation, cells were harvested and lysed by freeze-thawing, and the clarified lysates were used for two additional blind virus passages, during which RNA samples were extracted from these lysates for further verification of viral replication by RT-PCR using gs8-specific primers (Table S2). These recombinant viruses were verified by Sanger sequencing of the gs8-NSP2 RT-PCR products (GenBank IDs: MW074066, MW074067, MW074067).

## Authors’ contribution

J.B.K.B., K.B., L.V., E.G., A.C. and A.B. designed and carried out experiments, and analyzed data. J.B., K.B., A.C. and A.B. jointly wrote the manuscript. R.T., D.C.L., C.D. contributed novel analytical tools. J.B. collected and analyzed EM data. A.C. collected and analyzed HDX data. K.B. and A.B. collected and analysed single-molecule fluorescence data. A.B. managed the project. All authors contributed ideas, discussed the results and were involved in writing of the manuscript.

## Competing interests

The authors declare no competing interests.

## Data and materials availability

All data needed to evaluate the conclusions in the paper are present in the paper and/or the Supplementary Materials. Sequencing data are available for rescued recombinant rotaviruses (GenBank IDs: MW074066, MW074067, MW074067). Preliminary Full wwPDB EM Map/Model Validation reports are included into the submission.

## Acknowledgements

The authors would like to thank Prof Ben Luisi, Dr Anders Barth and Dr Chris Hill for their valuable comments and suggestions. We would like to thank Dr Guido Papa and Dr Oscar Burrone for their generous gift of pT7 constructs used for rescuing recombinant rotaviruses.

Funding sources: A.B. acknowledges support from a Sir Henry Dale Fellowship jointly funded by the Wellcome Trust and the Royal Society (Grant Number: 213437/Z/18/Z); Biotechnology and Biological Sciences Research Council (BBSRC) White Rose DTP [BB/M011151/1 to J.P.K.B.]; European Regional Development Fund [CZ.02.1.01/0.0/0.0/15_003/0000441 to R.T.]; A.N.C. acknowledges support from a Sir Henry Dale Fellowship jointly funded by the Wellcome Trust and the Royal Society (Grant Number 220628/Z/20/Z) and a University Academic Fellowship from the University of Leeds. E.H.G. holds a Biomedicine Discovery Scholarship and is an EMBL-Australia PhD student. C.D. is an EMBL-Australia Group Leader and acknowledges support from the ARC (DP190103407) and the NHMRC (APP1162921 & APP1184637).

Deutsche Forschungsgemeinschaft SFB1032 (Project B3) [to D.C.L.] and the Ludwig-Maximilians-Universität, München through the Center for NanoScience (CeNS) and the LMUinnovativ initiative BioImaging Network (BIN) (to D.C.L.).

Funding from the BBSRC (BB/M012573/1) to purchase HDX-MS instrumentation is acknowledged.

Funding for open access charge: Wellcome Trust.

## Supplementary Information

**Supplementary Figure S1.**
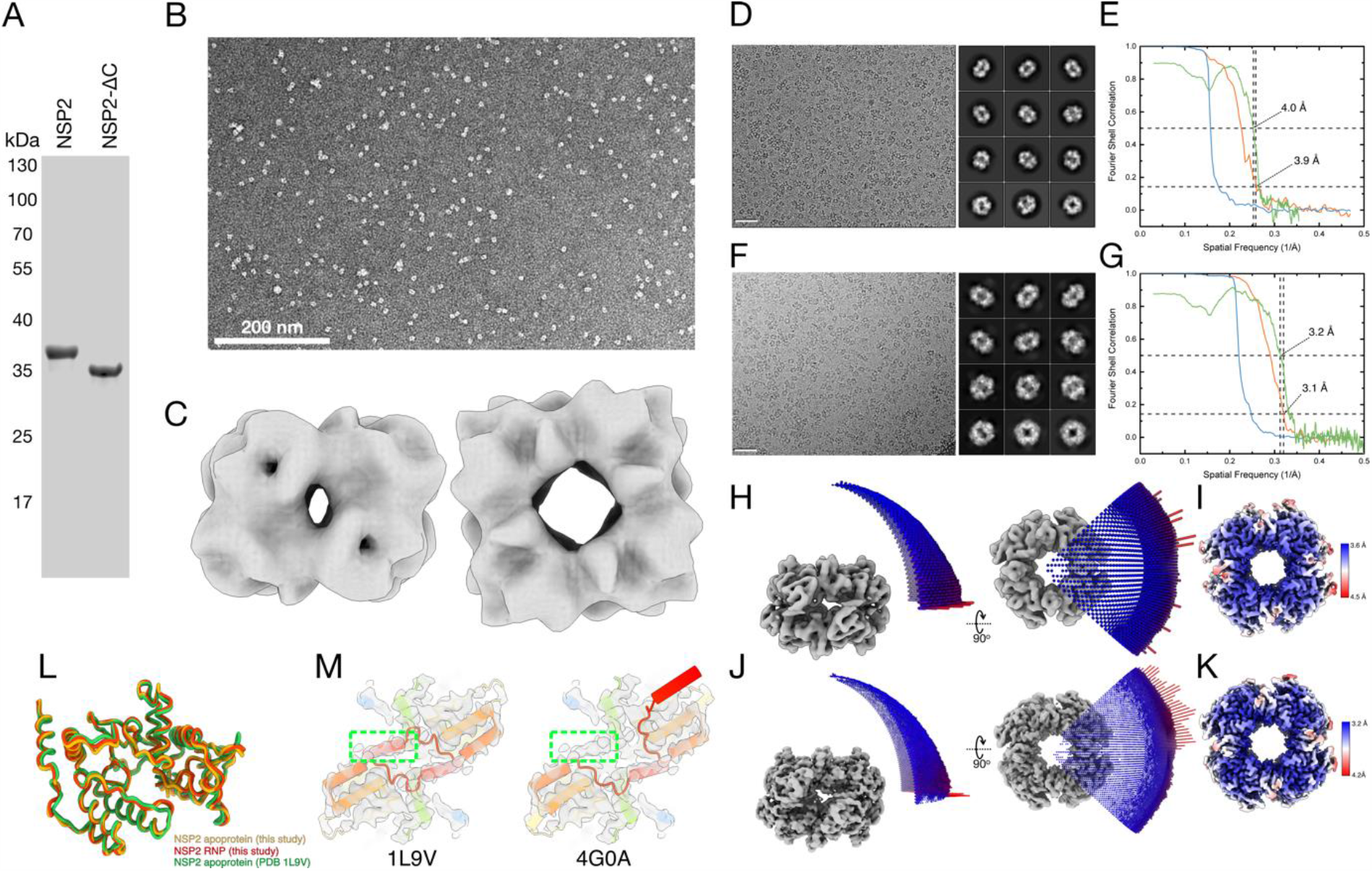
NSP2–ΔC variant assembles into octamers. **A:** SDS-PAGE of purified NSP2 and NSP2-ΔC. **B:** Representative negative stain EM micrograph of NSP2-ΔC **C:** 3D reconstruction of NSP2-ΔC. **D**: A representative cryo-EM micrograph of the NSP2 apoprotein and corresponding 2D class averages. **E:** The Fourier shell correlation (FSC) curves for the overall NSP2 apoprotein map (orange) and phase randomized map (blue). The map has a final resolution of 3.9 Å. Model versus map FSC curve (green) indicates a model resolution of 4.0 Å. **F:** A representative cryo-EM micrograph of the NSP2 RNP complex and corresponding 2D class averages. **G:** The Fourier shell correlation (FSC) curves for the overall NSP2-RNP complex map (orange) and phase randomized map (blue). Map has a final resolution of 3.2 Å. Model versus map FSC curve (green) indicates a model resolution of 3.1 Å. **H & J:** Euler angle distribution of particles corresponding to NSP2 apoprotein and NSP2-RNP reconstructions. The cylinder height and color represent the number of particles (blue to red – low to high). A D4 symmetry has been applied for both the NSP2 apoprotein and the NSP2-RNP complex reconstructions. **I & K:** NSP2 apoprotein (F) and NSP2-RNP complex (H) reconstructions colored by local resolution as calculated by Relion. **L:** Aligned models of the NSP2 apoprotein and the NSP2-RNP complex built from reconstructions presented in this study (yellow and red, respectively), and a previous crystal structure of the NSP2 apoprotein (PDB ID 1L9V) (green). Alignment of the apoprotein and RNP models from this study to 1L9V had an RMSD of 1.081 Å and 0.772 Å, respectively. The two models from this study had an RMSD of 0.742 Å. **M:** Fitting of previous NSP2 crystal structures into the NSP2 apoprotein cryo-EM density map. The “open” NSP2 conformation (PDB 4G0A) has the CTR flipped outwards, while the “closed” conformation (PDB 1L9V) has the CTR making contacts with the NSP2_core_. The open conformation is not represented by the cryo-EM map, as the CTH density is clearly visible and localized to the NSP2_core_. The green box highlights the cryo-EM density corresponding to the C-terminus.

**Supplementary Figure S2.**
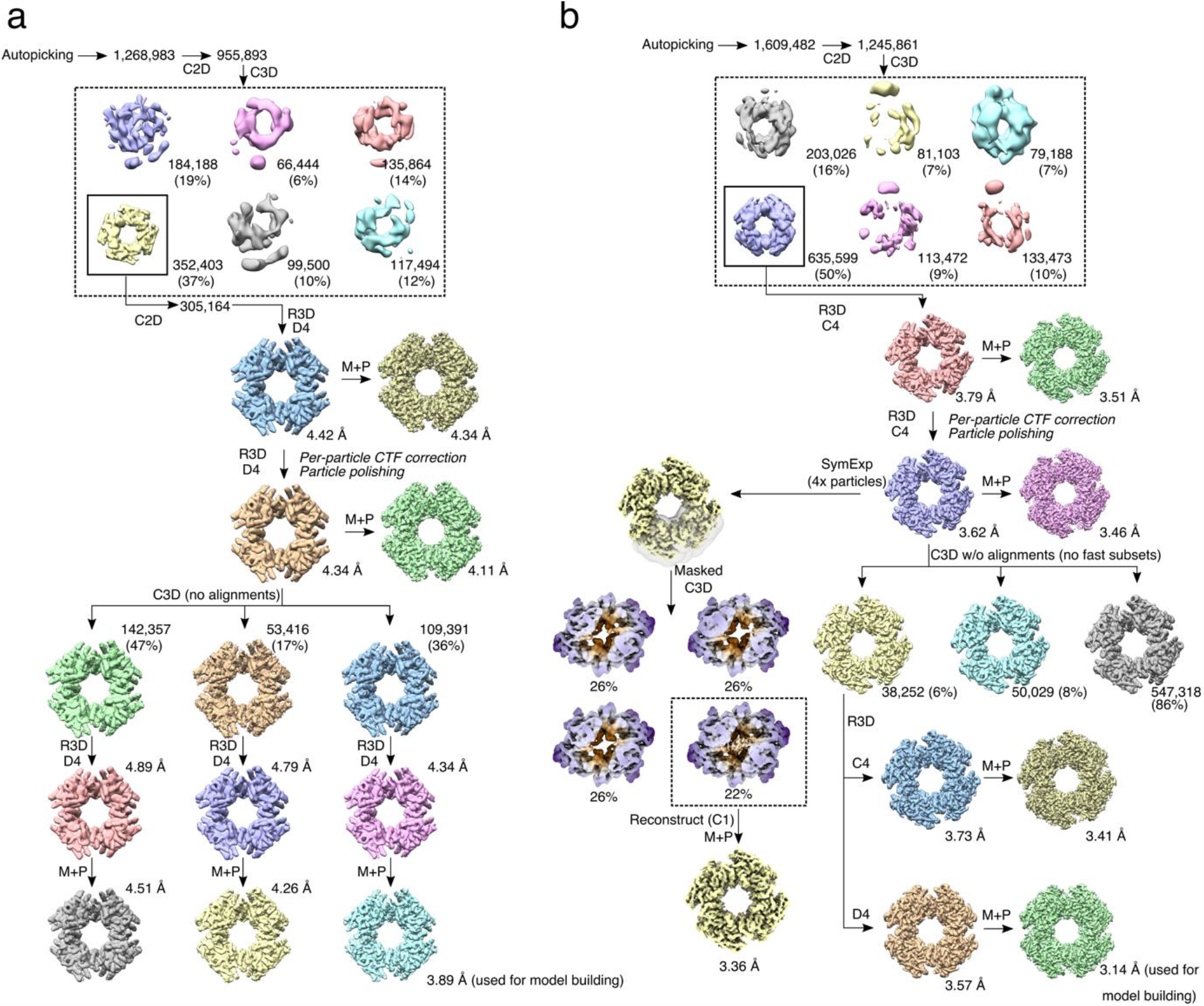
Cryo-EM structure determination of the NSP2 apoprotein and the NSP2-RNP complex. **A**: The image processing workflow used to determine the 3D reconstruction of the NSP2 apoprotein. Initial rounds of 2D classification (C2D) and 3D classification (C3D) were performed using Fast Subsets and image alignments with 25 iterations. The highest quality 3D class average was subjected to further 2D classification without fast subsets, and used for 3D refinement (R3D) with D4 symmetry imposed. After per-particle CTF correction and particle polishing, further 3D classification was performed without fast subsets, and without performing image alignment. 3D reconstructions were ultimately subjected to masking and post-processing (M + P). **B**: The image processing scheme used to determine the 3D reconstruction of NSP2 RNP. Initial 2D and 3D classifications were performed with fast subsets. Purple: Focused classification of NSP2-RNP demonstrate the existence of an RNA-binding groove with variable occupancy. Exemplary 3D class averages of NSP2-RNP with strong (boxed) and weak (all other) RNA density features.

**Supplementary Figure S3.**
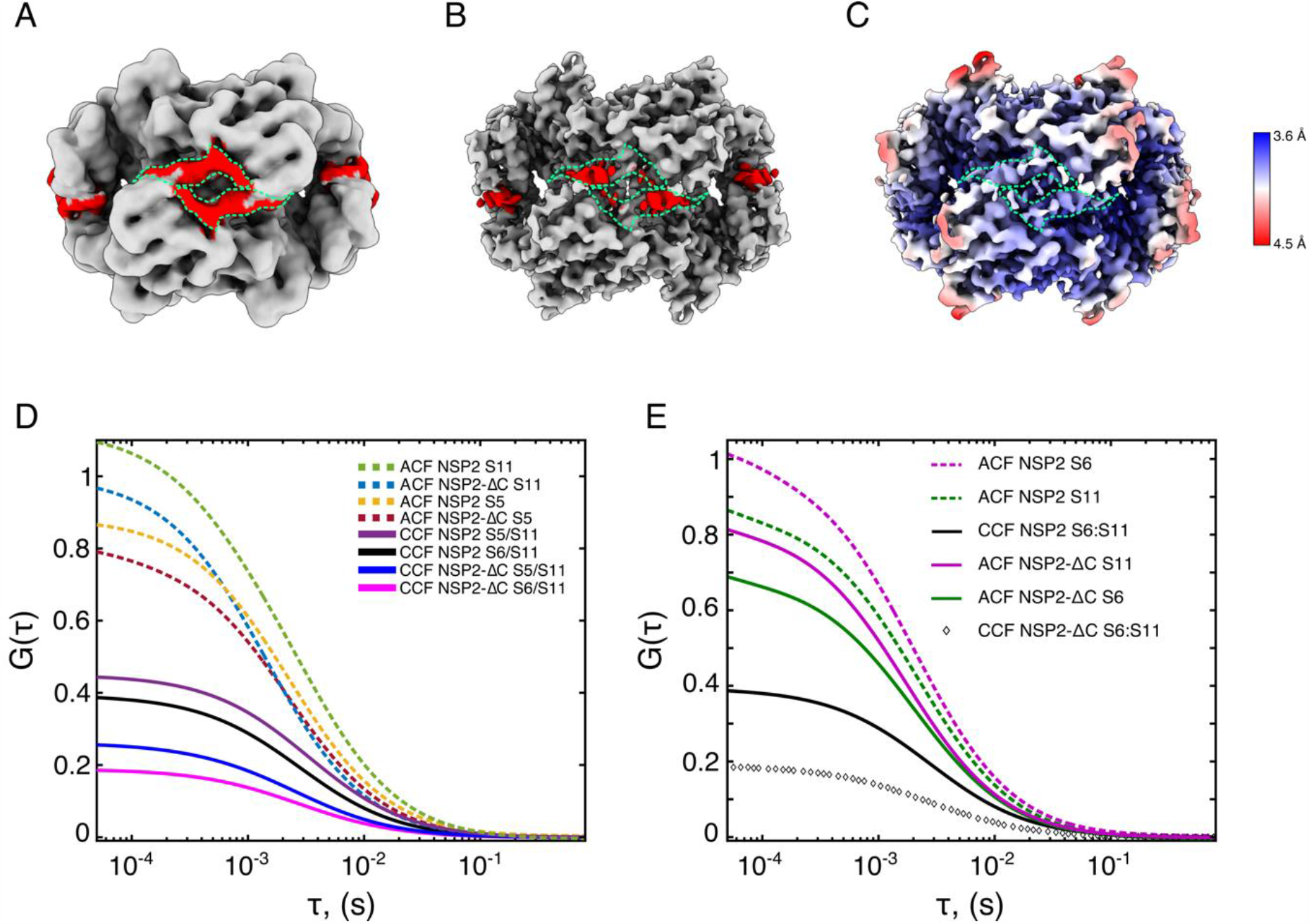
Role of the CTR in promoting RNA-RNA interactions. **A & B:** Unsharpened (A) and sharpened (B) NSP2 apoprotein cryo-EM density maps. Density corresponding to the CTR is shown in red (green outline). Linker density is diffuse in the sharpened map (B). **C:** Local resolution of NSP2 with the CTR outlined in green. **D:** Autocorrelation functions (ACFs) of rotavirus (RV) segments S5 and S11 in the presence of NSP2 and NSP2-ΔC (dashed lines). Cross-correlation functions (CCFs) (bold lines) of segments S5 and S11 in the presence of NSP2 (burgundy) and NSP2-ΔC (navy) and segments S6 and S11 in the presence of NSP2 (black) and NSP2-ΔC (magenta). **E:** ACFs and CCFs of S6 and S11 in the presence of NSP2 and NSP2-ΔC. Dashed, colored lines correspond to ACFs of WT NSP2 in complex with RV S6 and S11 RNAs, and continuous, colored lines correspond to ACFs of NSP2-ΔC in complex with RV S6 and S11 RNAs. Continuous black line and diamonds correspond to CCFs of both S6 and S11 in the presence of NSP2 and NSP2-ΔC, respectively.

**Supplementary Figure S4.**
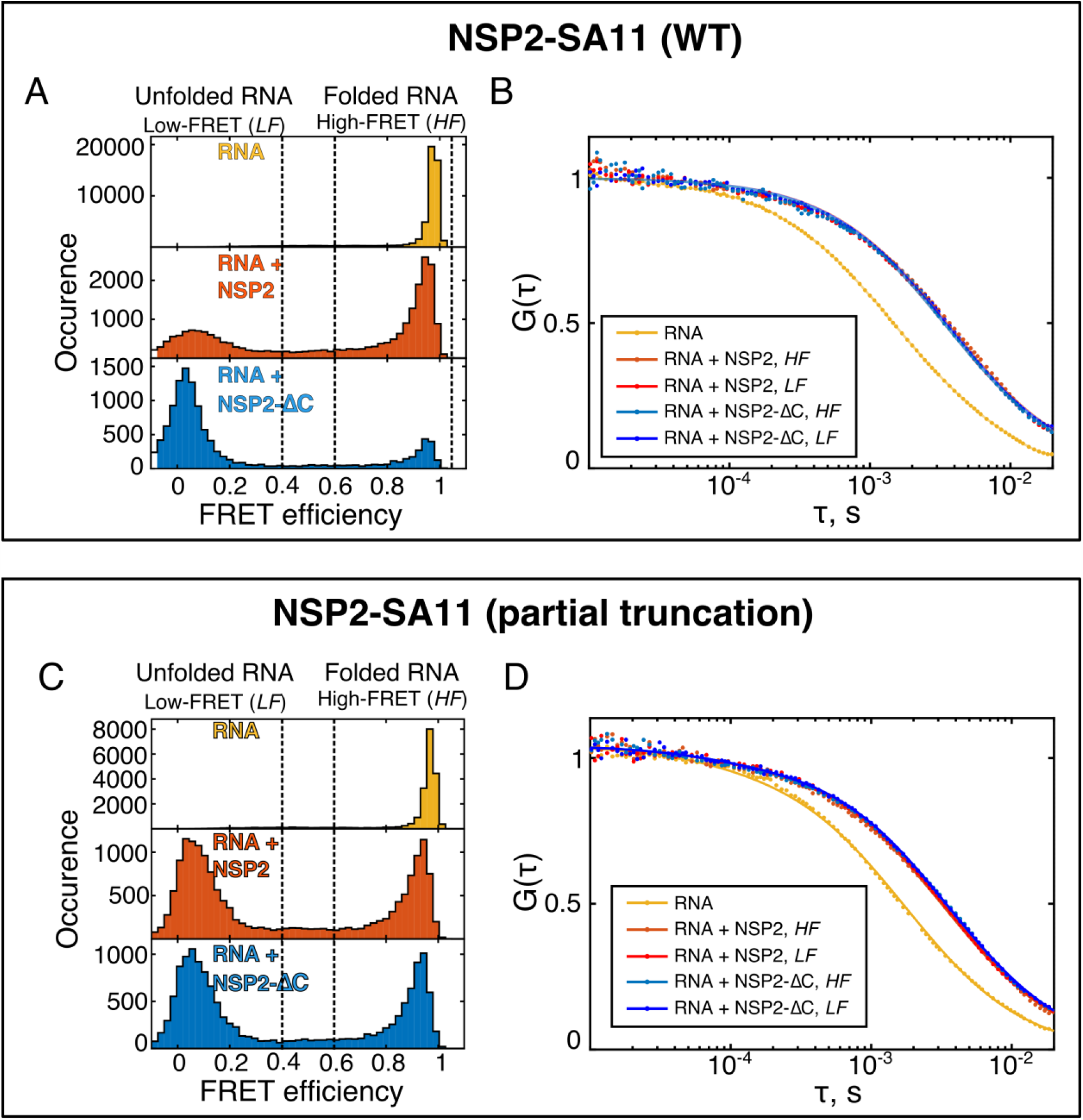
SmFRET and burstwise FCS analysis of stem-loop unwinding by NSP2 variants. **A & B:** SmFRET histograms (**A**) and corresponding burstwise FCS analysis (**B**) of the NSP2 RF strain and NSP2(RF)-ΔC. **C & D** smFRET histograms (**C**) and corresponding burstwise FCS analysis (**D**) of the NSP2 SA11 strain and a partially truncated NSP2 (lacking the unstructured residues 314-316). These residues are not essential for virus replication (*9*), and do not contribute to RNA unwinding.

**Supplementary Figure S5.**
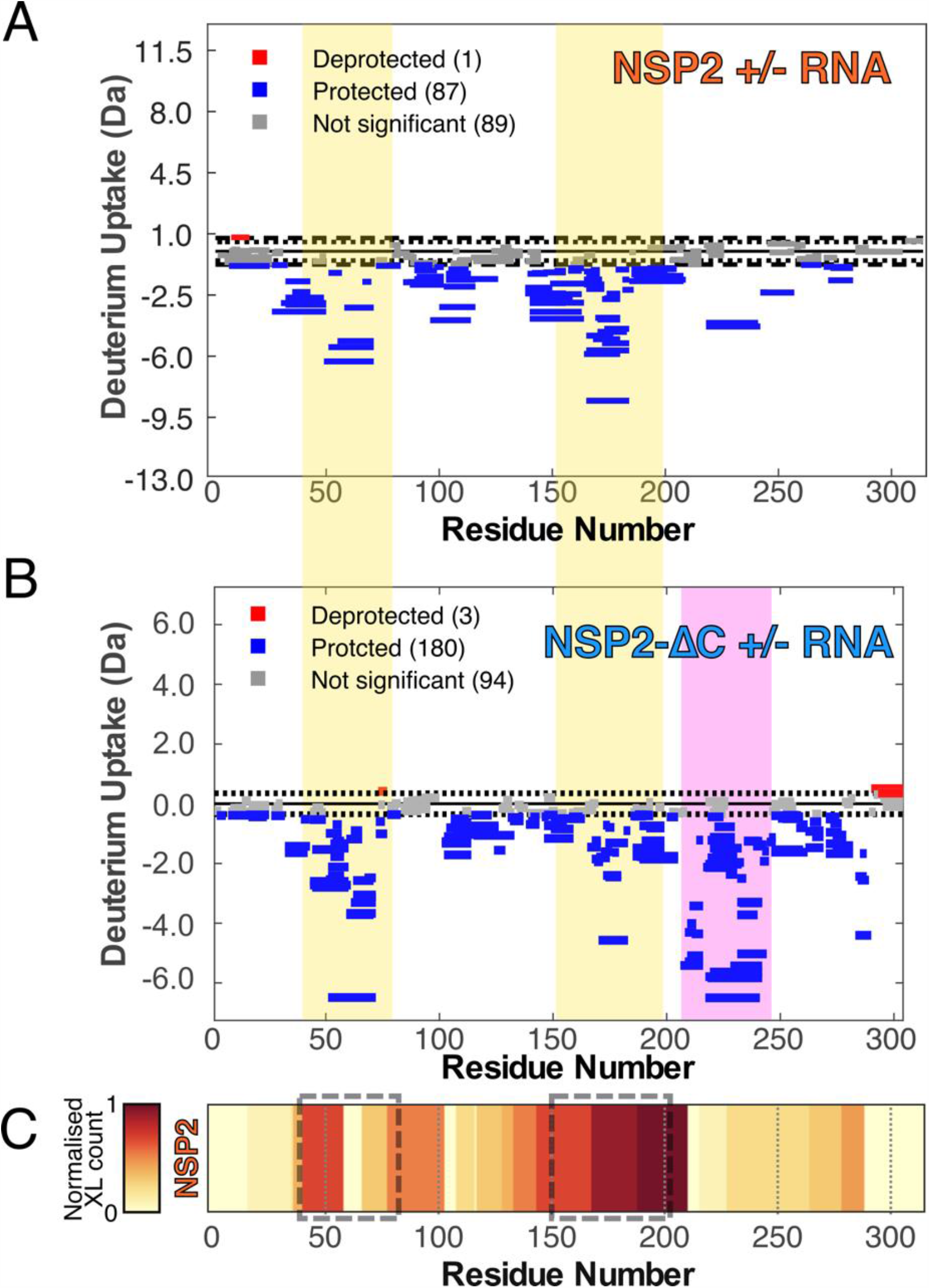
**A & B:** Woods plots showing summed differences in deuterium uptake in NSP2 **(A)** and NSP2-ΔC **(B)** over all timepoints, comparing the apoproteins with their respective RNP complex. Regions corresponding to flexible loops located within the polar, equatorial groove of NSP2 are denoted by yellow shading. In NSP2-ΔC, the magenta box denotes residues that would be otherwise buried underneath the CTR. This includes R240. **C**: A heatmap of crosslinked peptides mapped onto the NSP2 sequence (plotted as counts per residue). Residues corresponding to flexible loops within the equatorial grove are denoted by grey boxes. These regions correspond to the yellow-shaded areas in the above Woods plots.

**Supplementary Figure S6.**
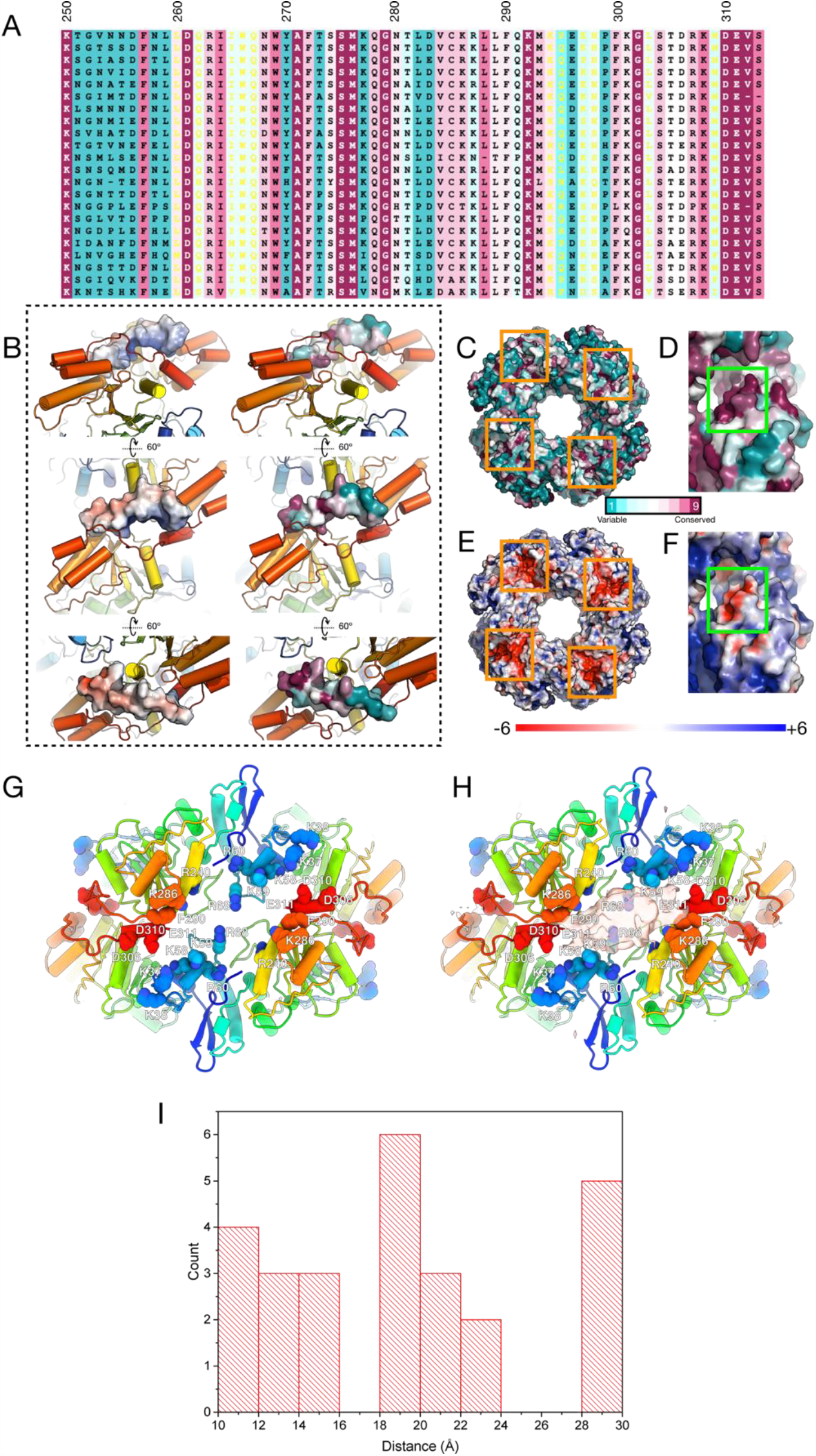
**A:** A ConSurf analysis of the NSP2 C-terminus (residues 250 – 313). **B:** Charge clusters patches are observable on alternate sides of the ampholytic NSP2 CTR surface. The cartoon representation of NSP2 is rainbow colored according to sequence position. A single CTR is shown as a molecular surface structure. Left: ABPS surface. Right: ConSurf surface conservation surface. **C – F:** Surface-exposed acidic patches within the NSP2 with low (C & E, orange boxes) and high (D & F, green boxes) levels of sequence conservation, as calculated using ConSurf. The highly conserved acidic patch in D & F is within the CTR (D306, D310, E311). **G:** An atomic model of NSP2 with potential RNA-interacting residues & acidic residues within CTR (D306, D310, E311) shown as spheres. **H:** The model shown in A with the positive cryo-EM difference map (i.e. RNA density) superimposed. **I**: Distances between nearest CTR and cryo-EM-identified RNA-binding residues (KKRR K58, K59, R60, R68) and acidic residues within the CTR (DDE D306, D310, E311). For the sake of clarity, only distances between RNA-binding residues and CTR 30 Å were included. Distances were measured using ChimeraX (*48*).

**Supplementary Figure S7.**
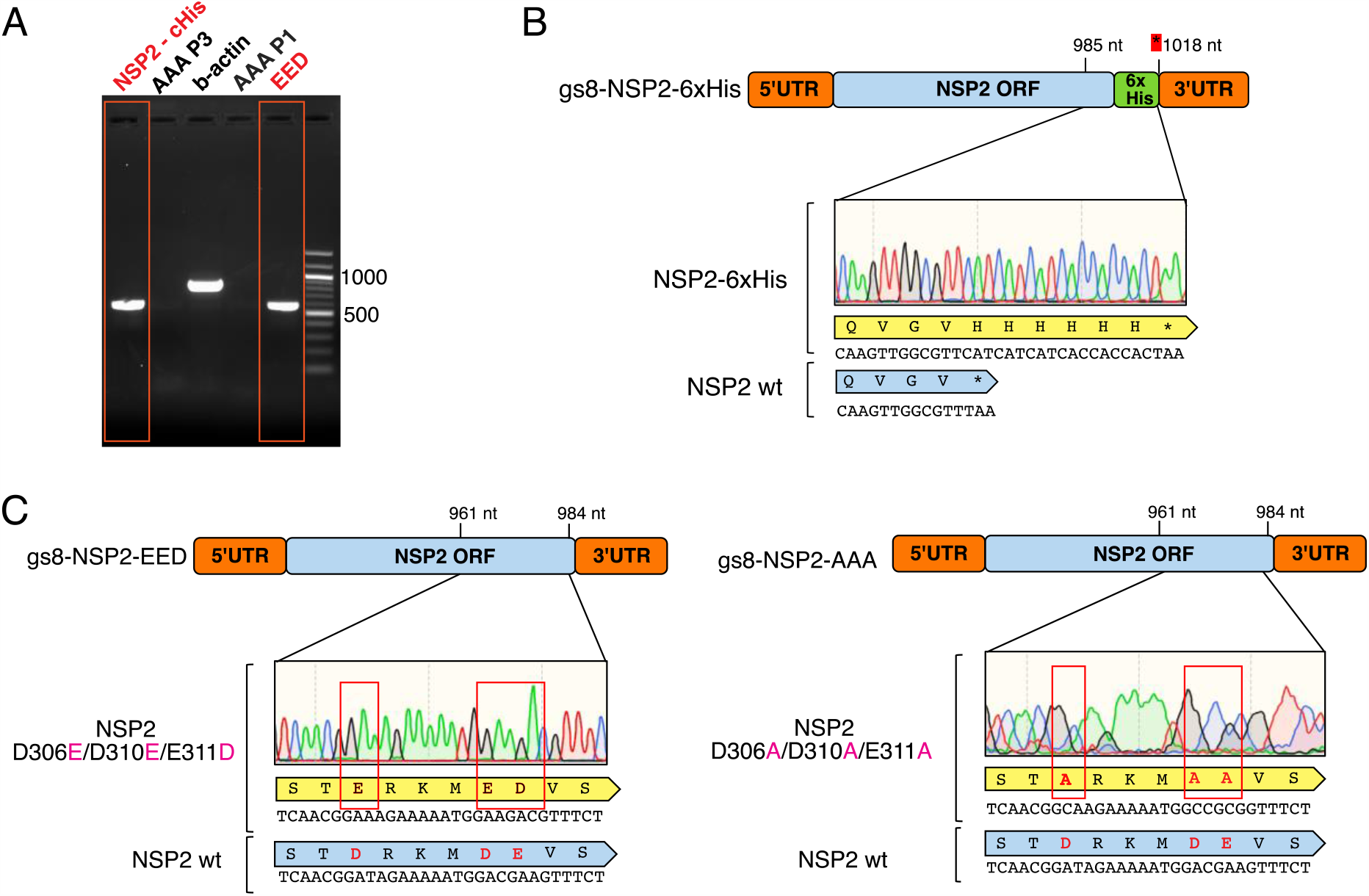
Rescue of replication-competent NSP2 mutant rotaviruses in reverse genetics experiments. A. RT-PCR results of the RNA extracted from MA104 cells infected with lysates from reverse genetics experiments designed to rescue C-terminally 6xHis-tagged NSP2 (panel B), and NSP2-EED and NSP2-AAA mutants (panel C). For each experiment, three independent attempts were made to rescue the wild-type virus and the mutants. Total RNA was extracted from virus-infected cells for each mutant (Materials and Methods), and amplified using gs8-specific primers, prior to further verification by sequencing. For gs8-NSP2-AAA mutant, sequencing data are shown only for the plasmid gs8-NSP2-AAA used for the virus rescue, as no cDNA could be made due to the absence of replicating viral RNA, even after two blind passages (panel A, AAA mutant, lane P3).

**Table S1.**
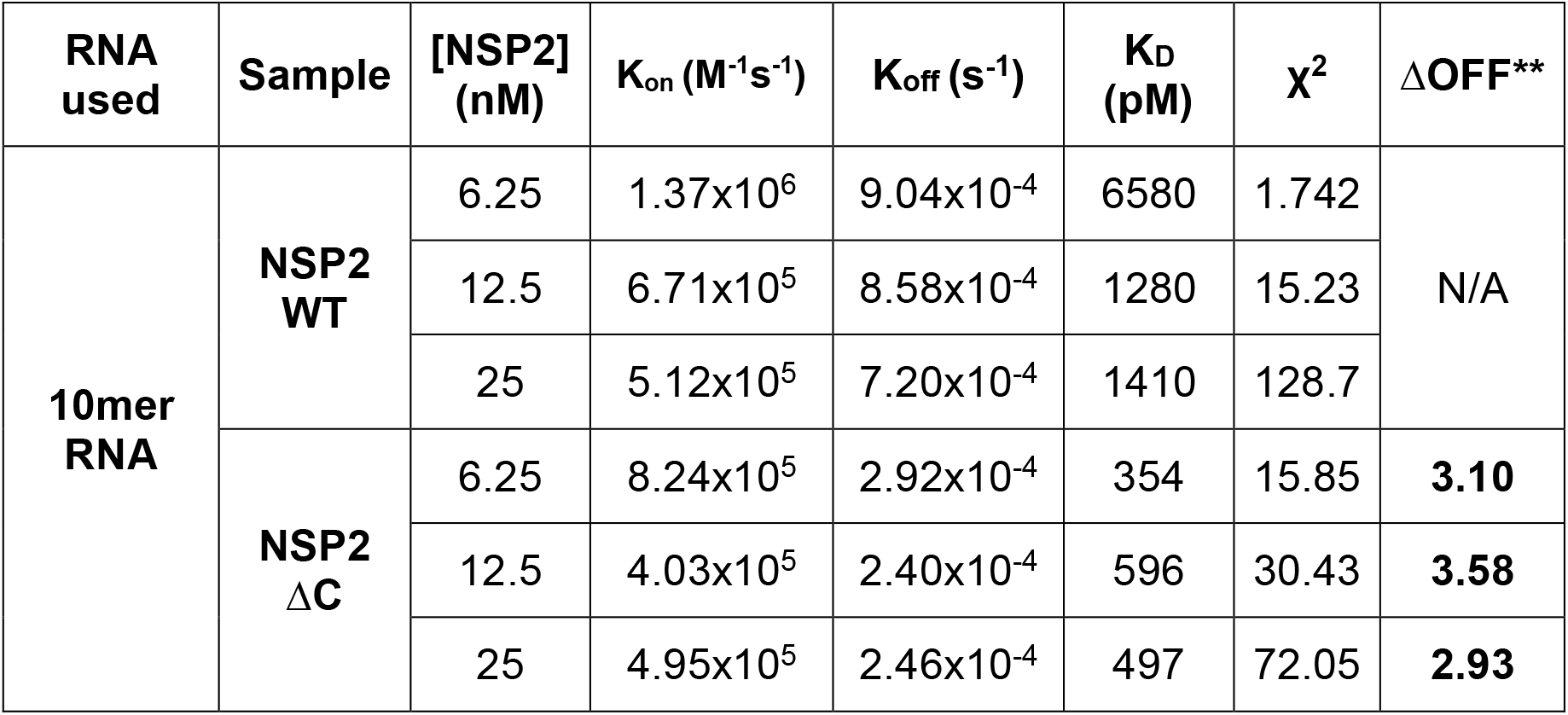
Binding kinetics of NSP2 and NSP2-ΔC as measured by SPR. The association (K_on_) and dissociation (K_off_) rate constants obtained by SPR are given below.

| RNA used | Sample | [NSP2] (nM) | $K_{on}$ ($M^{-1}s^{-1}$) | $K_{off}$ ($s^{-1}$) | $K_D$ (pM) | $\chi^2$ | $\Delta OFF^{**}$ |
| --- | --- | --- | --- | --- | --- | --- | --- |
| 10mer RNA | NSP2 WT | 6.25 | $1.37 \times 10^6$ | $9.04 \times 10^{-4}$ | 6580 | 1.742 | N/A |
| | | 12.5 | $6.71 \times 10^5$ | $8.58 \times 10^{-4}$ | 1280 | 15.23 | |
| | | 25 | $5.12 \times 10^5$ | $7.20 \times 10^{-4}$ | 1410 | 128.7 | |
| | NSP2 $\Delta C$ | 6.25 | $8.24 \times 10^5$ | $2.92 \times 10^{-4}$ | 354 | 15.85 | **3.10** |
| | | 12.5 | $4.03 \times 10^5$ | $2.40 \times 10^{-4}$ | 596 | 30.43 | **3.58** |
| | | 25 | $4.95 \times 10^5$ | $2.46 \times 10^{-4}$ | 497 | 72.05 | **2.93** |

**Table S2.**
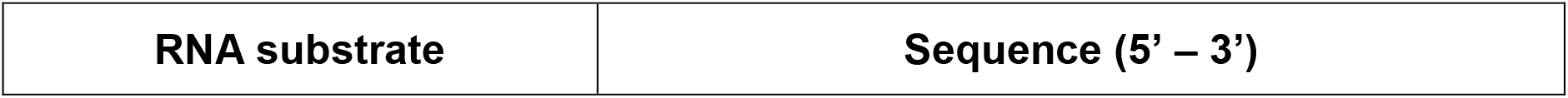
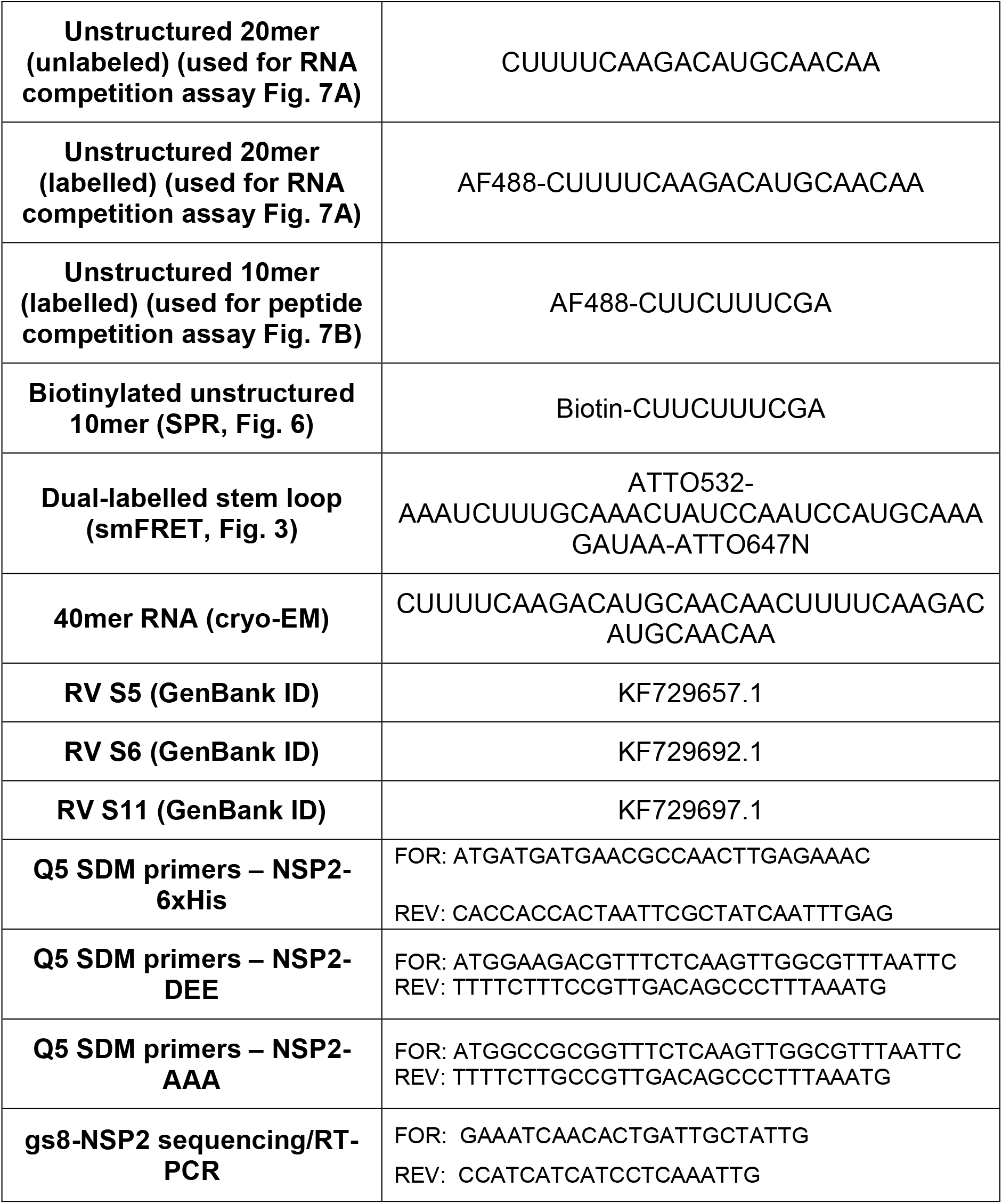
Sequences of RNAs used in the study

**Table S3.** Cryo-EM data collection and refinement statistic

|  | **NSP2 apoprotein** | **NSP2 RNP** |
| --- | --- | --- |

| **Data collection and processing** |  |  |
| --- | --- | --- |
| Nominal Magnification | 75,000 |  |
| Voltage (kV) | 300 kV |  |
| Defocus range (μm) | -1.2 to -2.4 | -1.5 to -3.5 |
| Detector | Falcon III | Gatan K2 |
| Pixel size (Å) | 1.065 | 1.07 |
| Electron exposure (e<sup>-</sup>/Å<sup>2</sup>) | 110 | 55 |
| Symmetry imposed | D4 |  |
| Particles in first 3D classification | 955,893 | 1,245,861 |
| Final particle number | 109,391 | 38,252 |
| Nominal map resolution (Å) | 3.90 | 3.14 |
| B-factor* | -239.2 | -100.8 |
| FSC threshold | 0.143 |  |
| Map resolution range (Å) | 3.64 – 4.46 | 3.22 – 4.19 |
| **Refinement, model composition and validation** |  |  |
| Initial model used | PDB 1L9V |  |
| Model resolution (Å) | 4.0 | 3.2 |
| FSC threshold | 0.5 |  |
| Non-hydrogen atoms | 20368 | 20368 |
| RMSD bonds (Å) | 0.004 | 0.008 |
| RMSD angles (°) | 0.651 | 0.810 |
| ***Ramachandran*** |  |  |
| Favoured (%) | 92.52 | 93.01 |
| Allowed (%) | 7.48 | 6.99 |
| Outliers (%) | 0.00 | 0.00 |
| Rotamer outliers (%) | 0.00 | 0.00 |
| Clash score | 10.77 | 7.13 |
| Molprobity score | 2.02 | 1.84 |
| ***Model-to-map fit*** |  |  |
| Cross-correlation coefficient (mask) | 0.82 | 0.83 |
| Cross-correlation coefficient (volume) | 0.81 | 0.81 |
| Main-chain | 0.81 | 0.83 |
| Sidechain | 0.78 | 0.81 |

## Notes

### Competing Interest Statement

The authors have declared no competing interest.

